# Screening the PRISM library against *Staphylococcus aureus* reveals a sesquiterpene lactone from *Liriodendron tulipifera* with inhibitory activity

**DOI:** 10.1101/2022.06.03.494747

**Authors:** Riley D. Kirk, Margaret E. Rosario, Nana Oblie, Terra Marie M. Jouaneh, Marina A. Carro, Christine Wu, Elizabeth Leibovitz, Elizabeth Sage Hunter, Robert Literman, Sara M. Handy, David C. Rowley, Matthew J. Bertin

## Abstract

Infections caused by the bacterium *Staphylococcus aureus* continue to pose threats to human health and put a financial burden on the healthcare system. The overuse of antibiotics has contributed to mutations leading to the emergence of methicillin-resistant *Staphylococcus aureus*, and there is a critical need for the discovery and development of new antibiotics to evade drug resistant bacteria. Medicinal plants have shown promise as sources of new small molecule therapeutics with potential uses against pathogenic infections. The Principal Rhode Island Secondary Metabolite (PRISM) library is a botanical extract library generated from specimens in the URI Heber W. Youngken Jr. Medicinal Garden by upper-division undergraduate students. PRISM extracts were screened for activity against strains of methicillin-susceptible *S. aureus* (MSSA). An extract generated from the tulip tree (*Liriodendron tulipifera*) demonstrated growth inhibition against MSSA, and a bioassay-guided approach identified a sesquiterpene lactone, laurenobiolide, as the active constituent. Intriguingly, its isomers tulipinolide and epi-tulipinolide lacked potent activity against MSSA. Laurenobiolide also proved to be more potent against MSSA than the structurally similar sesquiterpene lactones constunolide and dehydrocostus lactone. Laurenobioloide was most abundant in the twig bark of the tulip tree, supporting the historical and cultural usage of twig bark in poultices and teas.

**ABSTRACT GRAPHIC:** 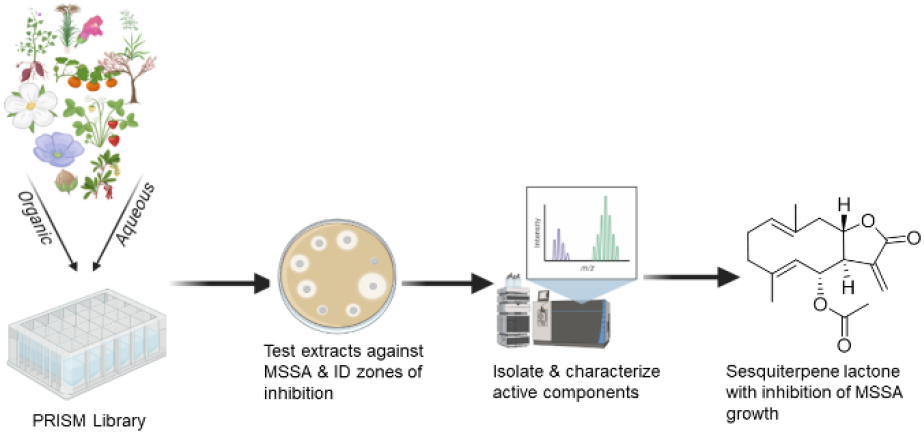

## INTRODUCTION

Infections caused by pathogenic bacteria claim millions of lives yearly and accrue billions of dollars in healthcare costs.^1^ The discovery and implementation of antibiotics to combat pathogens has saved millions of lives worldwide since the discovery of penicillin by Sir Alexander Fleming in 1928,^2,3^ Apart from inhibitors of cell wall biosynthesis like the β-lactam antibiotics, other classes of antibiotic molecules with distinct mechanisms of action such as the inhibition of protein biosynthesis, the inhibition of DNA synthesis, or the inhibition of folic acid metabolism are now routinely utilized in the clinic.^4^ The pharmacounderline>phores for the majority of these drugs were initially identified from natural products, primarily from microbial sources.^5^ However, the emergence and spread of antibiotic drug resistance among bacterial pathogens now represent an existential threat to our healthcare systems.^6^

*Staphylococcus aureus* causes severe health and economic problems associated with morbidity, mortality, and extended hospital stays due to invasive infections.^7^ While methicillin-resistant *S. aureus* (MRSA) is of major concern in hospital settings, methicillin-susceptible *S. aureus* (MSSA) can be the primary cause of invasive *S. aureus* infections in the hospital.^8^ Examining an infant cohort with invasive *S. aureus* infections from 1997-2012 showed that infant mortality after infection was similar for MRSA and MSSA, and MSSA actually caused more infant infections and more deaths than MRSA.^9^ Methicillin-susceptible *Staphylococcus aureus* (MSSA) can infect multiple human organs, including the skin which is the most prevalent. Untreated MSSA can lead to invasive infections in the bloodstream that lead to life threatening conditions such as sepsis.^10^

Plant metabolites offer promise in providing new molecular scaffolds to use as antibiotics either in isolation or in combination with currently used treatments to increase efficacy.^11,12^ The same specialized metabolites that a plant may use to combat microbes in the environment can potentially be harnessed to combat human pathogens. The tulip tree (*Liriodendron tulipifera)* has been of interest to medicinal chemists for its purported biological properties and cultural and historical use. *Liriodendron tulipifera* is a large flowering deciduous tree endemic across the eastern United States. Indigenous Americans valued the tulip tree for its timber. Medicinally it was utilized as a tonic, antipyretic, and anti-malarial agent.^13-15^ During the Civil War, Samuel Preston Moore, Surgeon General of the Confederacy, responded to the blockade of medicines by the Union Army by building new field hospitals and commissioning a study of medicinal plants of the Southeast to treat soldiers. Francis Porcher led the search for alternative plant medicines, conducting a deep study of the Southeast principally based on the cultural legacy of Cherokee healers. He ultimately compiled a book containing plant remedies native to the Southern States to help soldiers deal with injuries.^16^ One of the plants he highlighted was the tulip tree which was used for rheumatism, gout, laxative, headache, malaria, and other purposes.^16-18^ The described bioactivities were later validated by discovery of anti-plasmodial aporphine alkaloids and sesquiterpene lactones from the tree.^19^ While antibacterial activity has been demonstrated historically from plant preparations and recently from extracts,^15^ no studies have yet identified the antibacterial constituents of the tulip tree.

The Heber W. Youngken Jr. Medicinal Garden at the University of Rhode Island is a botanical collection of nearly 300 medicinal plant species. In 2019, we developed an extract library of aqueous and organic extracts of the aerial and root portions of specimens found in the garden.^20^ This resource, the Principal Rhode Island Secondary Metabolite (PRISM) library with 135 extracts at present was screened for antibiotic activity against MSSA.

Herein, we identified the organic (methanol) extract of the tulip tree leaves (*Liriodendron tulipera*) as having growth inhibitory properties against *S. aureus* DMS 1104, an MSSA strain. Using bioassay-guided isolation, the bioactive constituent was identified as the sesquiterpene lactone, laurenobiolide. We further investigated the relative abundance of this compound in various parts of the tulip tree demonstrating that the compound was present in higher concentrations in the twig bark when compared to other plant parts. Additionally, we examined other Magnoliaceae specimens to determine if they produced laurenobiolide and other related sesquiterpene lactones. Finally, we evaluated a panel of structurally similar sesquiterpene lactones (Figure 1) for their ability to inhibit MSSA.

**Figure 1.**
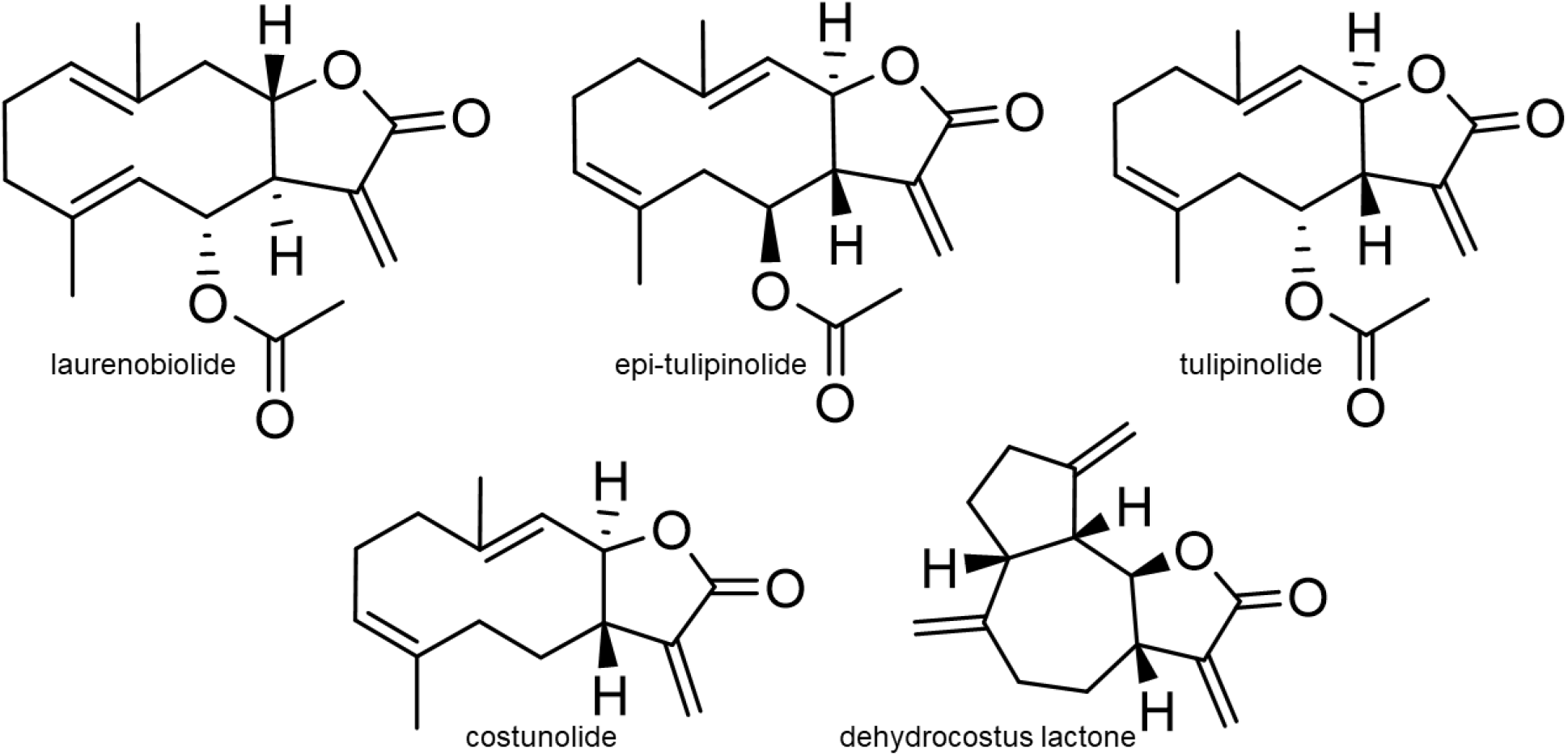
The primary active compound against MSSA was determined to be the sesquiterpene lactone laurenobiolide in this study. Other sesquiterpene lactones identified from *Liriodendron tulipifera* and magnolias were tulipinolide, epi-tulipinolide, costunolide, and dehydrocostus lactone.

## RESULTS AND DISCUSSION

Screening the PRISM library for anti-MSSA activity revealed that a methanol extract derived from *L. tulipifera* provided reproducible activity against methicillin-susceptible *S. aureus* (Figure S1). It was subsequently prioritized for further study due to its documented historical uses as an antimicrobial treatment. High performance liquid chromatography (HPLC) (Figure S2) was used to visualize the complex metabolite profile leading to our strategy for bioassay-guided fractionation. The tulip tree extract was first fractionated on a C18 column using a stepwise gradient of CH_3_OH in H_2_O. Five fractions were generated, and the most robust activity against MSSA was observed from constituents that eluted with 60% and 80% CH_3_OH (Figure S3). A more polar fraction also initially displayed antibacterial activity, but upon retesting individually isolated components, it seemed to be an additive effect of multiple compounds rather than a single potent entity responsible for the activity. Upon visual analysis of the HPLC chromatograms for each fraction, the most active fractions contained two large peaks that eluted at 30 minutes from the column (Figure S3). We speculated that a compound in one of these peaks could be responsible for the observed activity.

The two prominent peaks were isolated using semi-preparative HPLC and the latter eluting peak exhibited the largest zone of inhibition in the MSSA assay (Figure S4A). The peak purity was checked using a phenyl-hexyl column paired with HPLC-DAD analysis. We determined that the earlier eluting peak (peak 1) was a single pure compound while the later eluting peak was two separate analytes (Figure S4B). Upon repurification and retesting of the three compounds, it was determined that only peak 2.1 had antimicrobial activity, while peaks 1 and 2.2 did not show any observable activity in the disk diffusion assay (Figure S4C). Analysis and dereplication via 1D and 2D NMR (Figure S5 – S9 and S11) and HRESIMS (Figure S10) revealed that these three metabolites were isomers and the active peak 2.1 was determined to be laurenobiolide. HRESIMS of 2.1 gave an *m/z* of 313.1418 ([M+Na^+^], calcd: 313.1416 for C_17_H_22_O_4_Na). The specific rotation measured for peak 2.1: [α]^23^_D_ 16.8 (*c* 1.0, EtOH) closely matched the literature value of [α]^25^_D_ 17.1 (EtOH) for laurenobiolide.^21^ Additionally, examination of the ^1^H NMR spectra of peak 2.1 showed an envelope of olefinic protons from *δ* 4.70 to 5.50 consistent with the original characterization of this metabolite by Tada and Takeda in 1971,^22^ and the ^1^H NMR spectrum clearly showed the presence of conformational isomers, a phenomenon previously described for laurenobiolide during NMR analysis (Figure S6).^23,24^ Examination of correlations from COSY, HSQC, and HMBC NMR data confirmed the identification (Figures S7-S9). Examination of ^1^H NMR spectra identified peak 1 as epi-tulipinolide (Figure S5), and peak 2.2 as tulipinolide (Figure S11). Evidence included a multiplet resonance at *δ* 5.72 in peak 1 corresponding to H-8, which was in a cluster of resonances between *δ* 4.76 and 5.03 in peak 2.2. Furthermore, the acetyl methyl groups at *δ* 2.09 and 2.06 were consistent with those of tulipinolide and epi-tulipinolide, respectively, as detailed in the original isolation and elucidation work, as were the olefinic methyl resonances.^25^ Interestingly, among these three sesquiterpene lactones that are extremely similar in structure (Figure 1), only laurenobiolide displayed antibiotic activity towards *S. aureus* DMS 1104.

The minimum inhibitory concentration (MIC) for laurenobiolide and a panel of sesquiterpene lactones was determined via broth dilution assay. Laurenobiolide showed the strongest potency with an MIC of 7.8 μg/mL. Costunolide, one of the terpenoids devoid of an acetyl group, had an MIC value of 31 μg/mL. Epi-tulipinolide, tulipinolide and dehydrocostus lactone had MIC values of 62 μg/mL, 250 μg/mL, and 62 μg/mL respectively.

Laurenobiolide was evaluated for its cytotoxic effects to HaCaT cells, a commonly studied human keratinocyte cell line. The compound demonstrated cytotoxic effects at 10 μM showing 13% cell viability as compared to vehicle control (Figure S12). Epi-tulipinolide and tulipinolide showed less cytotoxic activity against the HaCaT cells at 10 μM with cell viability of 66% and 84%, respectively (Figure S12). Doses at 1 μM and lower did not show cytotoxicity. These observations are consistent with those of Dettweiler and coworkers, who showed that branch bark extracts of *L. tulipifera* possessed significant mammalian toxicity when tested against cells.^15^

Next, we investigated which parts of the tulip tree contained the highest concentrations of laurenobiolide. HPLC-DAD and LC-MS/MS analysis were performed to evaluate the presence of this constituent in the trunk bark, leaves, and twigs of the tulip tree. Laurenobiolide was most abundant in the twigs, and low amounts were detected in the leaves, but no laurenobiolide was detected in the trunk bark (Figure S13). These findings could be an artifact of the sampling approach where outer bark pieces were removed non-invasively so as to not damage the tree.

Further analysis showed the presence of laurenobiolide in the twig bark, but not the inner twig tissue (Figure S14). These observations are consistent withs some of the documented traditional Cherokee uses of the twig bark for medicine,^26^ and these data will now inform future investigations of plant parts while provoking intriguing questions about the ecological role of laurenobiolide. Additionally, both the twig itself and a poultice-like preparation made with twigs and jojoba oil showed zones of inhibition when tested against MSSA, while other plant parts tested did not show activity including a portion of ‘stripped twig’ with the outer covering removed (Figure S15). This helps connect the historic use of the twig bark poultices with the data on laurenobiolide discovered in this report.

We also examined several magnolia species for the presence of laurenobiolide and other sesquiterpene lactones. Laurenobiolide, epi-tulipinolide, and tulipinolide were only detected in *L. tulipifera* (Figure 2A and 2B). However, constunolide was detected in *L. tulipifera, Magnolia virginiana*, and *Magnolia acuminata* (Figure 2C and Figure S16), and dehydrocostus lactone was detected in *M. macrophylla* and *M. virginiana* (Figure S17). This was confirmed by comparison of the extracts to authentic standards of dehydrocostus lactone and constunolide. Additionally, constunolide and dehydrocostus lactone were identified as a library hits on the GNPS Social Molecular Networking website following library analysis (Figures S16 and S17).

**Figure 2.**
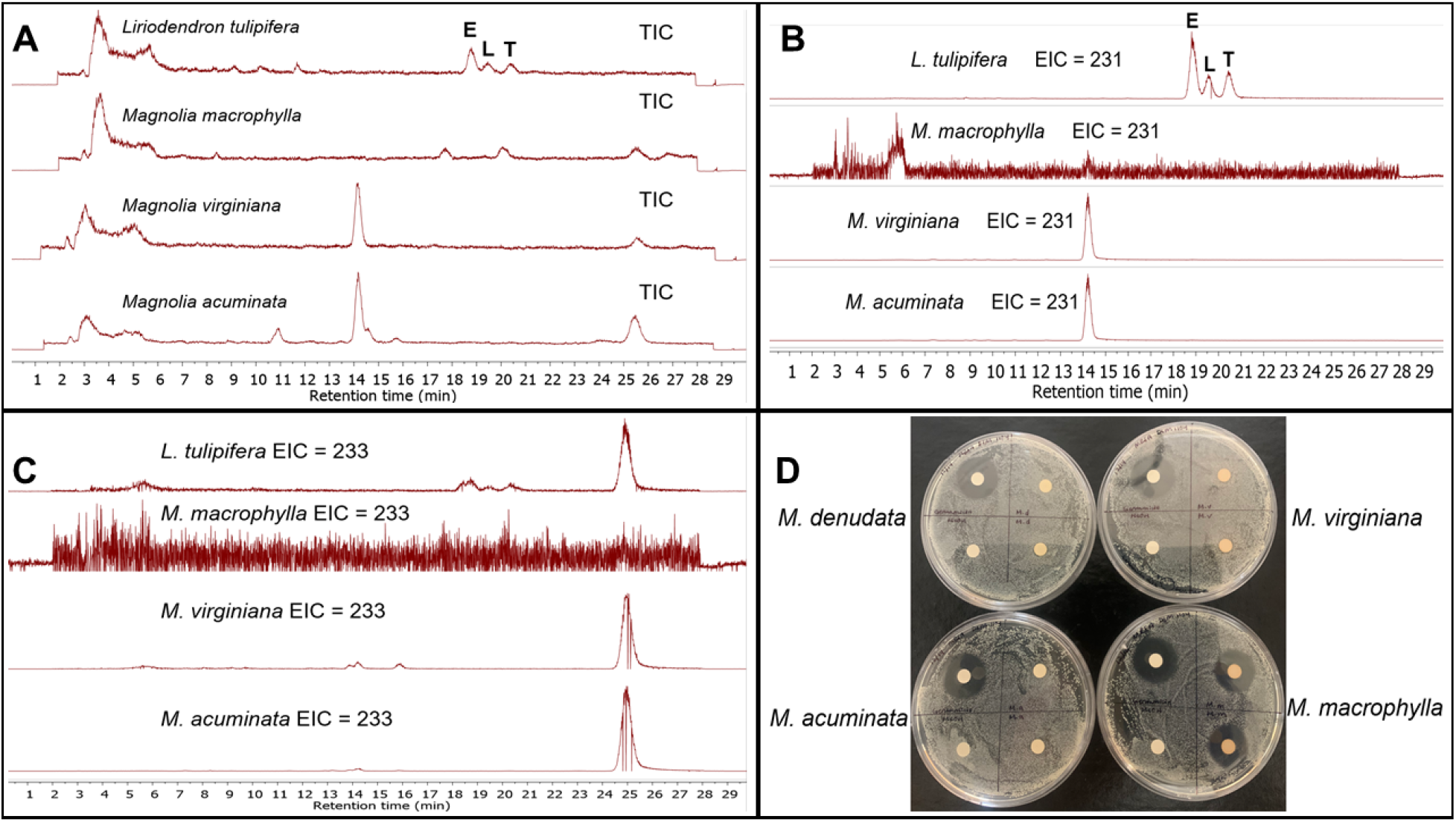
Sesquiterpene lactones in *Liriodendron tulipifera* and three other *Magnolia* spp. A) Epi-tulipinolide (E), laurenobiolide (L) and tulipinolide (T) were only detected in *L. tulipifera* extracts. B) An extracted ion chromatogram (*m/z* 231) shows the base peak of the three acetylated sesquiterpene lactones in *L. tulipifera*, and an unidentified metabolite in *M. virginiana* and *M. acuminata*. C) An extracted ion chromatogram (*m/z* 233) shows the presence of costunolide in *L. tulipifera, M. virginiana* and *M. acuminata*. D) Only the *M. macrophylla* extract showed anti-MSSA activity. Gentamicin sulfate, an aminoglycoside antibiotic, was used as the positive control (20 μg/disk; upper left quadrant on each plate). The test concentration of the extracts was 200 μg/disk, and each extract was tested in duplicate. Negative controls are in the bottom left quadrant of each plate.

Interestingly, other than *L. tulipifera*, only the *M. macrophylla* extract showed inhibition of MSSA amongst the magnolia extracts tested (Figure 2D). To determine the active components in the *M. macrophylla* extract, we conducted bioassay-guided isolation. Following additional inhibition tests, we characterized honokiol as the major antibacterial component in the *M. macrophylla* extract (MIC = 7.8 μg/mL against MSSA). Honokiol has reported potent anti-MSSA activity with MIC values as low as 2.5 μg/mL.^27^ The *M. denudata* extract did not show the presence of any of the known sesquiterpene lactones on which we focused (Figure S18) and its extract was inactive against MSSA (Figure 2D). Even though *M. virginiana* and *M. acuminata* extracts contained constunolide, which showed modest inhibitory activity against MSSA, the concentration was likely too low to show antibacterial activity.

While *Magnolia* and *Liriodendron* are both genera in the magnolia family, *Liriodendron* is the only genus in the Liriodendroidae subfamily, which consists of two species: *L. tulipifera* and *L. chinense*.^28^ Single nucleotide polymorphism analysis from DNA sequencing results of the specimen used in the current report clearly showed that the specimen was a *Liriodendron* and specifically *L. tulipifera* (Figure S19). It will be intriguing to examine authentic specimens of *L. chinense* to compare the metabolite profile and sesquiterpene lactone composition between the two *Liriodendron* species.

We searched for biosynthetic genes that could putatively generate sesquiterpene lactones such as laurenobiolide in publicly available *L. tulipifera* RNAseq data. The search was based on the pathway identified in *Helianthus annus*.^29^ This pathway synthesizes a sesquiterpene lactone from inunolide or eupatolide using germacrene-A synthase/hydroxylase, and a cytochrome P450 enzyme. This pathway is predicted to occur widely in Viridiplantae,^30^ and germacrene-A has been identified in the closely related genus *Magnolia*.^31^ We recovered *L. tulipifera* transcripts for farnesyl pyrophosphate synthase, cytochrome P450 (E-class, group IV), acetyl-CoA acetyltransferase (cytosolic), and multiple terpene cyclases including germacrene D synthase (Table S1). Although we found seven distinct terpene cyclases, we did not find a transcript with sufficient homology to indicate the presence of germacrene A synthase or germacrene-A hydroxylase in the *L. tulipifera* transcriptome. We would predict that laurenobiolide would derive from germacrene A acid. Terpene cyclases can have wide substrate affinity,^32^ and it is possible that laurenobiolide is being synthesized via a novel biosynthetic pathway. However, due to the lack of a reference genome (this analysis utilized RNAseq data) it is also possible these germacrene A genes were simply not expressed in the tissue at the time of collection. Further genomic and transcriptomic investigations are needed to discern whether this compound is being synthesized via an undescribed biosynthetic pathway, or with germacrene-A, as in *Helianthus annus*.

There has been a consistent effort to develop drugs with unique chemical scaffolds and new mechanisms of action to combat pathogen resistance.^33^ While relatively few antibacterial compounds have been identified in the terpene class, terpenes and terpenoids are an incredibly diverse group of phytochemicals that possess potential as potent antimicrobials.^34^ The exact mechanism of action of many terpene antibiotics is unknown, but Griffin and coworkers identified that certain terpene metabolites inhibit two essential processes, which include oxygen uptake and oxidative phosphorylation.^35^

## CONCLUSION

The tulip tree extract has documented antimicrobial activity,^15^ yet the constituents responsible for the activity had not been identified until now. In this report, we have determined for the first time that the metabolite laurenobiolide isolated from *Liriodendron tulipifera* possesses potent anti-MSSA activity and is the likely component responsible for the antiseptic activity ascribed to the tulip tree. This work sheds light on the intriguing story of the cultural and historical use of tulip tree preparations. The structural features that allow laurenobiolide to inhibit MSSA growth, while very closely related structures do not inhibit growth, remains an intriguing observation that we plan to explore in future studies. The most likely anti-MSSA structural features would appear to be the position of double bonds and/or the absolute configuration about the ring juncture. Furthermore, we have demonstrated the utility of the PRISM library as a source of molecular potential in therapeutic areas, encouraging its future expansion and use in screening efforts. Future research will focus on the mechanism of action of laurenobiolide and its ecological role for the tree. Additionally, we will continue to search for antibacterial components from magnolias and other botanical specimens.

## MATERIALS AND METHODS

### General Experimental Procedures

Optical rotations were measured using a Jasco P-2000 polarimeter. NMR spectra were recorded using an Agilent NMR 500 MHz spectrometer with (CD_3_)_2_SO (referenced to residual DMSO at *δ*_H_ 2.50 and *δ*_C_ 39.5) or CDCl_3_ (referenced to residual CHCl_3_ at *δ*_H_ 7.26 and *δ*_C_ 77.2) at 25 °C. HRESIMS analysis was performed using an AB SCIEX TripleTOF 4600 mass spectrometer with Analyst software. MS/MS data were recorded on this same instrument using the product ion function. LC-MS/MS was performed using a ThermoFisher LTQ XL mass spectrometer with an electrospray ionization (ESI) source, and certain mass spectrometry experiments (standard analysis) were completed using a ThermoFisher ISQ mass spectrometer. Both the LTQ and ISQ were coupled to Dionex Ultimate 3000 HPLC systems equipped with a micro vacuum degasser, an autosampler, and diode-array detector (DAD). Semi-preparative and analytical HPLC was carried out using a Dionex UltiMate 3000 HPLC system equipped with a micro vacuum degasser, an autosampler, and a DAD. Standard compounds, constunolide and dehydrocostus lactone, were purchased from Sigma-Aldrich.

### Plant Harvesting and Preparation of Plant Extracts

Specimens of *Liriodrendron tulipifera, Magnolia denudata, Magnolia virginiana, Magnolia acuminata*, and *Magnolia macrophylla* were harvested from the University of Rhode Island campus, and voucher specimens of the magnolias were deposited in the KIRI herbarium. Biological material from the *L. tulipifera* specimen is held in the lab of the corresponding author. Plant extracts in the PRISM library were prepared as previously described from specimens in the Heber W. Youngken Jr. Medicinal Garden.^20^ Briefly, plants were separated into aerial and underground portions, then further divided (twigs, leaves, etc.) for extraction in both aqueous and organic (methanol) solvents. Each solvent was vacuum filtered to remove particulates, then either freeze-dried or evaporated under reduced pressure to remove remaining solvent. Dried product was reconstituted at 10 mg/mL in DMSO or CH_3_OH depending on solubility.

### Methicillin-Susceptible *Staphylococcus aureus* Inhibition Assays

*S. aureus* strain DSM 1104 was cryopreserved and stored at -80 °C. The strain was grown in Luria Bertani (LB) broth (Difco, Fisher scientific, Waltham, MA) at 37 °C, 150 rpm for 20 h. Plant extracts were prepared as described above and 20 μL were loaded onto sterile antimicrobial susceptibility test disks (Oxoid, Thermo Fisher, Franklin, MA). The disks were air dried for 30 min. Control disks were prepared with 20 μL MeOH and similarly dried. The disks were then added to Petri dishes (100×15 mm) containing 15 mL of LB agar (Alpha biosciences, Baltimore, MD) inoculated with DSM 1104. Plant parts that were directly tested against DSM 1104 were first sterilized by wiping with CH_3_OH and then dried before testing. Gentamicin sulfate (20 μg/disk) was the positive control for MSSA. Negative vehicle controls were DMSO or CH_3_OH (20 μL). Plates were incubated at 37 °C for 18 h after which the diameters of the zones of growth inhibition were measured to the nearest mm.

### Minimum Inhibitory Concentration (MIC) of Sesquiterpene Lactones

Minimum inhibitory concentrations (MICs) were determined in duplicate by broth microdilution in LB medium as previously described,^36^ and in accordance with Clinical Laboratory Standards Institute (CLSI) standards.^37,38^ *Staphylococcus aureus* DSM 1104 was obtained from the American Type Culture Collection (ATCC 25923™). Test samples were prepared at 10 mg/mL in DMSO. Gentamicin sulfate (2 mg/mL) in H_2_O was the positive control and DMSO was the negative control.

### Bioassay Guided Fractionation of Crude Plant Extracts

The tulip tree leaf organic extract was further fractionated using a C18 SPE cartridge to rarify samples before additional testing. Five fractions were produced from the extract, beginning with the most polar (80% H_2_O: 20% CH_3_OH) and ending with the least polar (100% CH_3_OH). Following solvent removal, the chromatography fractions were reconstituted at 10 mg/mL in mobile phase followed by HPLC-DAD analysis. A 70 min gradient method was utilized beginning with 15% acetonitrile (CH_3_CN) for 10 min, then increasing the concentration of CH_3_CN until 45 min, with a 20 min hold at 100% CH_3_CN, followed by a return to starting conditions for 5 min. Chromatograms were visually evaluated in parallel with repeated antimicrobial assays to identify potentially active compounds.

### Isolation of Sesquiterpene Lactones and Honokiol from *Liriodendron tulipifera* and *Magnolia macrophylla*

Having identified the potential active components in the tulip tree chromatography fractions following initial chromatogram analysis, fractions 3 (60% H_2_O/40% CH_3_OH) and 4 (80% H_2_O/20% CH_3_OH) were combined and subjected to reversed-phase semi-preparative HPLC using a Kinetex 5 μm C18 column (250 × 10 mm) and a gradient method. The mobile phase was 60% CH_3_CN/40% H_2_O with 0.05% formic acid (FA) added to each solvent with a flow rate of 3.0 mL/min. Initial conditions were held for 10.5 min followed by a linear gradient to 12.0 min reaching 100% CH_3_CN and holding at this concentration for 4.0 min, ultimately returning to initial conditions from min 17-20. Two chromatographic peaks were collected at t_R_ 9.5 (peak 1 - epi-tulipinolide) and 10.0 min (peak 2), respectively. Peak 2 was further purified using a phenyl-hexyl 5 μm column (250 × 10 mm) and an isocratic method. The mobile phase was 55% CH_3_CN/45% H_2_O with 0.05% FA added to each solvent and a flow rate of 3 mL/min. Peaks 2.1 (laurenobiolide; t_R_ 13.5 min) and 2.2 (tulipinolide; t_R_ 14.25 min) were isolated. The CH_3_OH branch extract of *M. macrophylla* was subjected to RP-HPLC using the same isocratic method as described above. The mobile phase was 55% CH_3_CN/45% H_2_O with 0.05% FA added to each solvent and a flow rate of 3 mL/min. A single peak eluting at 18.25 min was collected and identified as honokiol (4 mg).

### LC-MS/MS Analysis of *L. tulipifera* Plant Parts and Magnolia Specimens and Database Searching

Samples of tulip tree trunk, twig, and leaf were taken from a *L. tulipifera* specimen on the URI campus and methanol extracts were made using 1.0 g portions of each *L. tulipifera* plant part. Additionally twig extracts were generated from 1 g portions of *M. denudata, M. virginiana, M. acuminata*, and *M. macrophylla*. Each extract was subjected separately to HPLC-DAD and LC-MS/MS analysis with a specific scan event recording MS/MS spectra in data-dependent acquisition mode. A Kinetex 5 μm C18 column (150 × 4.6 mm) was used for separation of analytes in the extracts. The LC method consisted of a linear gradient from 15% to 100% CH_3_CN in H_2_O + 0.1% FA over 20 min, followed by an isocratic period at 100% CH_3_CN of 5 min. The flow rate was held constant at 0.4 mL/min. The MS spray voltage was 3.5 kV with a capillary temperature of 325 °C. For the MS/MS component, the CID isolation width was 1.0 and the collision energy was 35.0 eV. Following acquisition, the raw data files were converted to .mgf format using MSConvert from the ProteoWizard suite (http://proteowizard.sourceforge.net/tools.shtml). Library searching was performed using the online platform at Global Natural Products Social Molecular Networking website (gnps.ucsd.edu).^39^

### Detection of Sesquiterpene Lactones in *L. tulipifera* Plant Parts and Magnolias

*L. tulipifera* twig, leaf, and trunk bark CH_3_OH extracts (10 mg/mL) were subjected to HPLC-DAD analysis to evaluate the presence of laurenobiolide. Branch CH_3_OH extracts from *L. tulipifera, M. denudata, M. virginiana, M. acuminata*, and *M. macrophylla* were subjected to HPLC-DAD and LC-MS/MS analysis to specifically investigate the presence of the sesquiterpene lactones epi-tulipinolide, laurenobiolide, and tulipinolide. The chromatography parameters for HPLC-DAD and LC-MS/MS were identical. The mobile phase was 55% CH_3_CN/45% H_2_O with 0.05% formic acid added to each solvent and flow rate of 0.6 mL/min. Following the HPLC-DAD and LC-MS/MS analysis, the presence of costunolide and dehydrocostus lactone was confirmed by chromatographic comparison to authentic standards using the method described above.

### Culturing Human Keratinocytes (HaCaT)

Human keratinocytes (HaCaT) were a kind gift from the laboratory of Dr. Navindra Seeram at URI, and originally purchased from American Type Cell Culture Collection. The culture was maintained in Dulbecco’s modified Eagle’s medium (DMEM) and the DMEM was supplemented with 10% fetal bovine serum. The incubator was kept at 37 °C with 5% CO_2_ at constant humidity. All test samples were dissolved in sterile DMSO for viability assessments.

### Cytotoxic Evaluation of Keratinocytes (HaCaT) in the Presence of *L. tulipera* Compounds

The viability of HaCaT cells in the presence of laurenobiolide, tulipinolide, and epi-tulipinolide was evaluated using cell titer glo (CTG 2.0). HaCaT cells were seeded in a sterile 96-well plate at 5×10^3^ cells per well and incubated for 12 h. Post incubation, the media was removed, and compounds were added along with new media, followed by a 24 h incubation. The CTG 2.0 reagent was added to each well and shaken at 200 rpm for 2 min using an orbital shaker. The plate was allowed to equilibrate at room temperature for 10 min, and then the luminescence was recorded using a Spectramax M2 plate reader. Viability was determined by comparing luminescence values of treated wells to those of the vehicle control (DMSO) (n = 4 per treatment). Cannabidiol (100 μM) was used as a positive control.^40^

### Identifying Putatively Useful Nuclear Markers from Whole-Genome Sequence (WGS) Data

We used the SISRS bioinformatics package^41^ to develop nuclear genetic markers for the (1) delineation of *Liriodendron* spp. from other Magnoliaceae and (2) to delineate *L. chinese* and *L. tulipifera*. WGS data for six *L. tulipifera*, six *L. chinese*, and 83 specimens of *Magnolia* spp. were downloaded from the NCBI SRA database (https://www.ncbi.nlm.nih.gov/sra). All read data were trimmed using the *bbmap* suite (https://sourceforge.net/projects/bbmap/) using a sliding-window with a Q10 cutoff and removing reads with a final average quality less than Q15 and/or length less than 50 bp. To enrich each dataset for nuclear data, we then used *bbmap* to remove any reads that mapped against a pooled set of 253 chloroplast assemblies and 5 mitochondrial assemblies downloaded from NCBI. Based on a nuclear genome size estimate for the group of 2.5 Gb, we subset reads from each species such that the final combined read depth was ∼25 Gb (10X coverage), with reads sampled evenly across the 69 species and specimens therein. These subset reads were then pooled and used as input for a genome assembly using Ray with default parameters,^42^ with the resulting assembly representing loci that are conserved enough among species to be assembled and compared easily among taxa.

### Identifying Fixed Sites for Use in Species and Genus Delineation Assays

To identify genomic sites that would be useful for species delineation between *Liriodendron* species, we mapped the trimmed reads from all twelve *Liriodendron* specimen against the SISRS composite genome using *bbmap* and identified all sites that were (1) fixed and identical within species for all six *L. tulipifera* or *L. chinese* specimens and (2) variable between species. A total of 164,710 species-specific SNPs were identified.

To identify SNPs that can delineate *Liriodendron* from *Magnolia*, we first identified all sites in the composite genome that were fixed and identical among all twelve *Liriodendron* specimens. Next, after setting aside eight *Magnolia* specimens for downstream testing, we removed any sites where those fixed *Liriodendron* alleles were found in any of the remaining 73 *Magnolia* samples. *Magnolia* samples had variable read coverage, and not finding a *Liriodendron* specific allele could be due to either (1) the allele not being present or (2) the allele not being sampled. To make a more conservate estimate of robust genus-specific SNPs, we only considered SNPs where there was read coverage in at least half (n=38) of the *Magnolia* specimens. A total of 630,741 *Liriodendron*-specific SNPs were identified.

### Testing Genus- and Species-Specific Genetic Markers

To assess the ability of the genetic SNP markers to identify samples to the genus and species level, we mapped read data from eight *Magnolia* specimens, two *L. tulipifera* samples, and one *L. chinese* sample against the SISRS composite genome. Alleles at sites identified above were queried for all samples, and the proportion of sites matching either (1) the *Liriodendron* genus-specific allele or (2) the *L. tulipifera* or *L. chinese* specific allele was tabulated.

For genus-level screening, individual samples had coverage for 83,908 – 594,416 genus SNPs per specimen. The three *Liriodendron* samples had the matching *Liriodendron* allele at 99.2% - 99.9% of surveyed sites, while no more than 0.44% of sites had the corresponding allele across the eight *Magnolia* samples. For species-level screening, the three *Liriodendron* samples had coverage for 22,094 – 152,985 sites per specimen. The *L. chinese*-specific allele was detected at over 99% of sites in the *L. chinese* sample, while fewer than 2% of sites contained the allele in either *L. tulipifera* sample. In parallel, 97.5% - 99.4% of sites in the two *L. tulipifera* samples had the *L. tulipifera*-specific allele, while it was found in only 0.21% of sites for the *L. chinese* sample.

### Evaluation of Potential Biosynthetic Genes

Publicly available *Liriodendron tulipifera* RNAseq data originally collected from apical shoots was retrieved from SRA (SRR16546781, SRR16546782, SRR16546783, SRR16546784, SRR16546785, SRR16546803, SRR16546804), trimmed with Trimmomatic (v0.39),^43^ and assembled using rnaSPAdes (v3.15.3).^44^ This resulted in 332,056 transcripts, with an average length of 1,011 bp. Transdecoder (https://github.com/TransDecoder/TransDecoder) was then run to predict reading frames of the resulting transcripts, resulting in 189,683 predicted proteins. These proteins were annotated with KEGG,^45^ Interpro,^46^ and a custom BLAST^47^ database comprised of biosynthetic genes that produced sesquiterpenes such as costunolide in other plant species.

## ASSOCIATED CONTENT

### Supporting Information

A supporting information document is available with this report and contains: HPLC chromatograms of tulip tree extracts and chromatography fractions and bioassay-guided isolation information, NMR and MS data for epi-tulipinolide, laurenobiolide, tulipinolide, and honokiol, cytotoxicity data for sesquiterpene lactones to HaCaT cells, and chromatography data on distribution of metabolite in plant parts and magnolia specimens.

Additionally, information on sequence analysis and putative biosynthetic enzymes are included.

## AUTHOR INFORMATION

### Author Contributions

Conceptualization, M.J.B., R.D.K. E.L.; methodology, R.D.K., M.E.R, N.O., T.M.J., M.A.C., C.W., and M.J.B; formal analysis, R.D.K., M.E.R., E.S.H, R.L., S.M.H., D.C.R., and M.J.B.; writing—original draft preparation, R.D.K., M.J.B.; writing—review and editing, R.D.K., M.E.R., T.M.J., E.L., D.C.R., and M.J.B. All authors have read and agreed to the published version of the manuscript.

### Notes

The authors declare no financial conflicts of interest.

## ACKNOWLEDGEMENTS

We would like to recognize that historical knowledge of *Liriodendron tulipifera* was the result of the efforts of indigenous people and their use of the tulip tree in traditional medicines. The University of Rhode Island itself occupies the traditional stomping ground of the Narragansett Nation and the Niantic People. We honor and respect the enduring and continuing relationship between the Indigenous people and this land by teaching and learning more about their history and present-day communities, and by becoming stewards of the land we, now too, inhabit. We thank Dr. Navindra Seeram at URI and his coworkers for the HaCaT cells. The abstract graphic was created with Biorender.com. In this study, the acquisition of high-resolution mass spectrometry, polarimetry, UV-vis data, and NMR data in this publication was made possible by the use of spectrometric and spectroscopic equipment and services available through the RIINBRE Centralized Research Core Facility, which is supported by the Institutional Development Award (IDeA) Network for Biomedical Research Excellence from the National Institute of General Medical Sciences of the National Institutes of Health under Grant P20GM103430. We additionally gratefully acknowledge research support for N.O. from the American Society of Pharmacognosy Summer Research Fellowship. The graphical abstract was created with BioRender.com.

## REFERENCES

(1) Dadgostar, P. Antimicrobial Resistance: Implications and Costs. Infect. Drug Rests. 2019, 12, 3903–3910.

(2) Sengupta, S.; Chattopadhyay, M. K.; Grossart, H. P. The multifaceted roles of antibiotics and antibiotic resistance in nature. Front. Microbiol. 2013, 4, 47.

(3) Ventola, C. L. The antibiotic resistance crisis: part 1: causes and threats. Pharm. Ther. 2015, 40(4), 277–283.

(4) Kapoor, G.; Saigal, S.; Elongavan, A. Action and resistance mechanisms of antibiotics: a guide for clinicians. J. Anaesthesiol. Clin. Pharmacol. 2017, 33(3), 300–305.

(5) Newman, D. J.; Cragg, G. M. Natural products as sources of new drugs over the nearly four decades from 01/1981 to 09/2019. J. Nat. Prod. 2020, 83(3), 770–803.

(6) Martens, E.; Demain, A. L. The antibiotic resistance crisis, with a focus on the United States. J. Antibiot. 2017, 70(5), 520–526.

(7) Chatterjee, A.; Rai, S.; Guddattu, V.; Mukhopadhyay, C.; Saravu, K. Is methicillin-resistant Staphylococcus aureus infection associated with higher mortality and morbidity in hospitalized patients? A cohort study of 551 patients from South Western India. Risk Manag. Healthc. Policy 2018, 11, 243–250.

(8) Crandall, H.; Kapusta, A.; Killpack, J.; Heyrend, C.; Nilsson, K.; Dickey, M.; Daly, J. A.; Ampofo, K.; Pavia, A. T.; Mulvey, M. A.; Yandell, M.; Hulten, K. G.; Blaschke, A. J. Staphylococcus aureus infection in Utah children; continued dominance of MSSA over MRSA. PLoS ONE 2020, 15(9), e0238991.

(9) Ericson, J. E.; Popoola, V. O.; Smith, P. B.; Benjamin, D. K.; Fowler, V. G.; Benjamin Jr., D. K.; Clark, R. H.; Milstone, A. M. Burden of invasive Staphylococcus aureus infections in hospitalized infants. JAMA Pediatr. 2015, 169(12), 1105–1111.

(10) van Hal, S. J.; Jensen, S. O.; Vaska, V. L.; Espedido, B. A.; Paterson, D. L.; Gosbell, I. B. Predictors of mortality in Staphylcoccus aureus bacteremia. Clin. Microbiol. Rev. 2012, 25(2):362–386.

(11) Fazly, B. S.; Khameneh. B.; Zahedian, M. R.; Hosseinzadeh H. In vitro evaluation of antibacterial activity of verbascoside, lemon verbena extract and caffeine in combination with gentamicin against drug-resistant Staphylococcus aureus and Escherichia coli clinical isolates. Avicenna J. Phytomed. 2018, 8(3), 246–253.

(12) Rossiter, S. E.; Fletcher, M. H.; Wuest, W. M. Natural products as platforms to overcome antibiotic resistance. Chem Rev. 2017, 117(19), 12415–74.

(13) Kang, Y. F.; Liu, C. M.; Kao, C. L.; Chen, C. Y. Antioxidant and anticancer constituents from the leaves of Liriodendron tulipifera. Molecules. 2014, 19, 4234–4245.

(14) Quassinti, L.; Maggi, F.; Ortolani, F.; Lupidi, G.; Petrelli, D.; Vitali, L. A.; Miano, A.; Bramucci, M. Exploring new applications of tulip tree (Liriodendron tulipera L.) Leaf essential oil as apoptic agent for human glioblastoma. Environ. Sci. Pollut. Res. 2019, 26, 30485–30497.

(15) Dettweiler, M.; Lyles, J. T.; Nelson, K. Dale, B.; Reddinger, R. M. Zurawski, D. V.; Quave, C. American Civil War plant medicines inhibit growth, biofilm formation, and quorum sensing by multidrug-resistant bacteria. Sci Rep. 2019, 9, 7692.

(16) Porcher, F. P. Resources of the southern fields and forests, medical, economical, and agricultural: being also a medical botany of the Confederate States, with practical information on the properties of the trees, plants, and shrubs. Steam-power press of Evans and Cogswell: Charleston, SC, 1863.

(17) Spencer, C. F.; Koniuszy, F. R.; Rogers, E. F.; Shavel, J.; Easton, N. R.; Kaczka, E. A.; Kuehl, F. A.; Phillips, R. F.; Walti, A.; Folkers, K.; Malanga, C.; Seeler, A. O. Survey of plants for antimalarial activity. Lloydia 1947, 10,145–174.

(18) Thacher, J. American medical biography; or, memoirs of eminent physicians who have flourished in America, to which is prefixed a succinct history of medical science in the United States from the First Settlement of the Country. 1967. Milford House, New York.

(19) Graziose, R.; Rathinasabapathy, T.; Lategan, C.; Poulev, A.; Smith, P. J.; Grace, M.; Lila, M. A.; Raskin, I. Antiplasmodial activity of aporphine alkaloids and sesquiterpene lactones from Liriodendron tulipifera L. J. Ethnopharmacol. 2011, 133(1), 26–30.

(20) Kirk, R. D.; Carro, M. A.; Wu, C.; Jamal, M.; Wharton, A. M.; Goldstein, D. G.; Rosario, M. E.; Gallucci, G. M.; Zhao Y.; Leibovitz E.; Bertin, M. J. Integrating natural product chemistry workflows into medicinal chemistry laboratory training: building the PRISM library and cultivating independent research. J. Chem. Educ. 2020, 98(2), 410–415.

(21) Doskotch, R. W.; Wilton, J. H.; Harraz, F. M.; Fairchild, E. H.; Huang, C.-T.; El-Feraly, F. S. Six additional sesquiterpene lactones from Liriodendron tulipifera. J. Nat. Prod. 1983, 46(6), 923–929.

(22) Tada, H.; Takeda, K. Structure of the sesquiterpene lactone laurenobiolide. J. Chem. Soc. D 1971, 21, 1391–1392.

(23) Tori, K.; Horibe, I.; Kuriyama, K.; Tada, H.; Takeda, K. Conformational isomers of laurenobiolide, a new ten-membered-ring sesquiterpene lactone. J. Chem. Soc. D 1971, 21, 1393–1394.

(24) Tori, K.; Horibe, I.; Tamura, Y.; Kuriyama, K.; Tada, H.; Takeda, K. Re-investigation of the conformation of laurenobiolide, a ten-membered ring sesquiterpene lactone by variable-temperature carbon-13 NMR spectroscopy. Evidence for the presence of four conformational isomers in solution. Tetrahedron Lett. 1976, 17(5), 387–390.

(25) Doskotch, R. W.; el Feraly, F. S. The structure of tulipinolide and epitulipinolide. Cytotoxic sesquiterpenes from Liriodendron tulipifera L. J. Org. Chem. 1970, 35(6), 1928–1936.

(26) Moerman, D. E. Native American ethnobotany. Timber Press Inc., Portland, OR, 1998.

(27) Choi, E. -J.; Kim, H. -I.; Kim, J. -A.; Jun, S. Y.; Kang, S. H.; Park, D. J.; Son, S. -J.; Kim, Y.; Shin, O. S. The herbal-derived honokiol and magnolol enhances immune response to infection with methicillin-sensitive Staphylococcus aureus (MSSA) and methicillin-resistant S. aureus (MRSA). Appl. Microbiol. Biotechnol. 2015, 99(10), 4387–4396.

(28) Chen, J.; Hao, Z.; Guang, X.; Zhao, C.; Wang, P.; Xue, L.; Zhu, Q.; Yang, L.; Sheng, Y.; Zhou, Y.; Xu, H.; Xie, H.; Long, X.; Zhang, J.; Wang, Z.; Shi, M.; Lu, Y.; Liu, S.; Guan, L.; Zhu, Q.; Yang, L.; Ge, S.; Cheng, T.; Laux, T.; Gao, Q.; Peng, Y.; Liu, N.; Yang, S.; Shi, J. Liriodendron genome sheds light on angiosperm phylogeny and species–pair differentiation. Nat. Plants 2019, 5 (1), 18–25.

(29) Frey, M.; Schmauder, K.; Pateraki, I.; Spring, O. Biosynthesis of eupatolide ? a metabolic route for sesquiterpene lactone formation involving the P450 enzyme CYP71DD6. ACS Chem. Biol. 2018, 13, 1536–1543.

(30) Hawkins, C.; Ginzburg, D.; Zhao, K.; Dwyer, W.; Xue, B.; Xu, A.; Rice, S.; Cole, B.; Paley, S.; Karp, P.; Rhee, S. Y. Plant Metabolic Network 15: A resource of genome-wide metabolism databases for 126 plants and algae. J. Integr. Plant Biol. 2021, 63, 1888–1905.

(31) Farag, M. A.; El Din, R. S.; Fahmy, S. Headspace analysis of volatile compounds coupled to chemometrics in leaves from the Magnoliaceae family. Rec. Nat. Prod. 2015, 9, 153–158.

(32) Pazouki, L.; Memari, H. R.; Kännaste, A.; Bichele, R.; Niinemets, Ü. Germacrene A synthase in yarrow (Achillea millefolium) is an enzyme with mixed substrate specificity: gene cloning, functional characterization and expression analysis. Front. Plant Sci. 2015, 6.

(33) Coates, A.R.M.; Halls, G.; Hu, Yanmin. Novel classes of antibiotics or more of the same? Br. J. Pharmacol. 2011, 163(1), 184–194.

(34) Mahizan, N.A.; Yang, S.; Moo, C.; Song, A.A.; Chong, C.; Chong, C.; Abushelaibi, A.; Lim, S.E.; Lai, K. Terpene Derivatives as a potential agent against antimicrobial resistance (AMR) Pathogens. Molecules 2019, 24(14), 2631.

(35) Griffin, S.G.; Wyllie, S.G.; Markham, J.L.; Leach, D.N.; The role of structure and molecular properties of terpenoids in determining their antimicrobial activity. Flavour Fragr. J. 1999, 14, 322–332.

(36) Deering, R. W.; Whalen, K. E.; Alvarez, I.; Daffinee, K.; Beganovic, M.; LaPlante, K. L.; Kishore, S.; Zhao, S.; Cezairliyan, B.; Yu, S.; Rosario, M.; Mincer, T. J.; Rowley, D. C. Identification of a bacteria-produced benzisoxazole with antibiotic activity against a multi-drug resistant Acinetobacter baumannii. J. Antibiot. 2021, 74, 370–380.

(37) Clinical and Laboratory Standards Institute. Methods for Dilution of Antimicrobial Susceptibility Tests for Bacteria that Grow Aerobically; Approved Standard ? 10th Edition. CLSI Document M07-A10. Clinical and Laboratory Standards Institute, Wayne, PA, 2015.

(38) Clinical and Laboratory Standards Institute. Performance Standards for Antimicrobial Susceptibility Testing; Twenty-Fifth Informational Supplement. CLSI Document M100-S25. Clinical and Laboratory Standards Institute, Wayne, PA, 2015.

(39) Wang, M.; Carver, J. J.; Phelan, V. V.; Sanchez, L. M.; Garg, N.; Peng, Y.; Nguyen, D. D.; Watrous, J.; Kapono, C. A.; Luzzatto-Knaan, T.; Porto, C.; Bouslimani, A.; Melnik, A. V.; Meehan, M. J.; Liu, W.-T.; Crüsemann, M.; Boudreau, P. D.; Esquenazi, E.; Sandoval-Calderón, M.; Kersten, R. D.; Pace, L. A.; Quinn, R. A.; Duncan, K. R.; Hsu, C.-C.; Floros, D. J.; Gavilan, R. G.; Kleigrewe, K.; Northen, T.; Dutton, R. J.; Parrot, D.; Carlson, E. E.; Aigle, B.; Michelsen, C. F.; Jelsbak, L.; Sohlenkamp, C.; Pevzner, P.; Edlund, A.; McLean, J.; Piel, J.; Murphy, B. T.; Gerwick, L.; Liaw, C.-C.; Yang, Y.-L.; Humpf, H.-U.; Maansson, M.; Keyzers, R. A.; Sims, A. C.; Johnson, A. R.; Sidebottom, A. M.; Sedio, B. E.; Klitgaard, A.; Larson, C. B.; Boya P C. A.; Torres-Mendoza, D.; Gonzalez, D. J.; Silva, D. B.; Marques, L. M.; Demarque, D. P.; Pociute, E.; O’Neill, E. C.; Briand, E.; Helfrich, E. J. N.; Granatosky, E. A.; Glukhov, E.; Ryffel, F.; Houson, H.; Mohimani, H.; Kharbush, J. J.; Zeng, Y.; Vorholt, J. A.; Kurita, K. L.; Charusanti, P.; McPhail, K. L.; Nielsen, K. F.; Vuong, L.; Elfeki, M.; Traxler, M. F.; Engene, N.; Koyama, N.; Vining, O. B.; Baric, R.; Silva, R. R.; Mascuch, S. J.; Tomasi, S.; Jenkins, S.; Macherla, V.; Hoffman, T.; Agarwal, V.; Williams, P. G.; Dai, J.; Neupane, R.; Gurr, J.; Rodríguez, A. M. C.; Lamsa, A.; Zhang, C.; Dorrestein, K.; Duggan, B. M.; Almaliti, J.; Allard, P.-M.; Phapale, P.; Nothias, L.-F.; Alexandrov, T.; Litaudon, M.; Wolfender, J.-L.; Kyle, J. E.; Metz, T. O.; Peryea, T.; Nguyen, D.-T.; VanLeer, D.; Shinn, P.; Jadhav, A.; Müller, R.; Waters, K. M.; Shi, W.; Liu, X.; Zhang, L.; Knight, R.; Jensen, P. R.; Palsson, B. Ø.; Pogliano, K.; Linington, R. G.; Gutiérrez, M.; Lopes, N. P.; Gerwick, W. H.; Moore, B. S.; Dorrestein, P. C.; Bandeira, N. Sharing and Community Curation of Mass Spectrometry Data with Global Natural Products Social Molecular Networking. Nat. Biotechnol. 2016, 34(8), 828–837.

(40) Olivas-Aguirre, M.; Torres-López, L.; Valle-Reyes, J. S.; Hernádez-Cruz, A.; Pottosin, I.; Dobrovinskaya, O. Cannabidiol directly targets mitochondria and disturbs calcium homeostasis in acute lymphoblastic leukemia. Cell Death and Dis. 2019, 10, 779.

(41) Schwartz, R. S.; Harkins, K. M.; Stone, A. C. Cartwright, R. A. A composite genome approach to identify phylogenetically informative data from next-generation sequencing. BMC Bioinformatics 2015, 16, 193.

(42) Boisvert, S.; Laviolette, F.; Corbeil, J. Ray: Simultaneous assembly of reads from a mix of high-throughput sequencing technologies. J. Comp. Biol. 2010, 17, 1519–1533.

(43) Bolger, A. M.; Lohse, M.; Usadel, B. Trimmomatic: a flexible trimmer for Illumina sequence data. Bioinformatics 2014, 30, 2114–2120.

(44) Bushmanova, E.; Antipov, D.; Lapidus, A.; Prjibelski, A. D. rnaSPADES: a de novo transcriptome assembler and its application to RNA-Seq data. GigaScience 2019, 8, giz100.

(45) Kanehisa, M.; Goto, S. KEGG: Kyoto encyclopedia of genes and genomes. Nucleic Acids Res. 2000, 28, 27–30.

(46) Blum, M.; Chang, H. -Y.; Chuguransky, S.; Grego, T.; Kandasaamy, S.; Mitchell, A.; Nuka, G.; Paysan-Lafosse, T.; Qureshi, M.; Raj, S.; Richardson, L.; Salazar, G. A.; Williams, L.; Bork, P.; Bridge, A.; Gough, J.; Haft, D. H.; Letunic, I.; Marchler-Bauer, A.; Mi, H.; Natale, D. A.; Necci, M.; Orengo, C. A.; Pandurangan, A. P.; Rivoire, C.; Sigrist, C. J. A.; Sillitoe, I.; Thanki, N.; Thomas, P. D.; Tosatto, S. C. E.; Wu, C. H.; Bateman, A.; Finn, R. D. The InterPro protein families and domains database: 20 years on. Nucleic Acids Res. 2021, 49, D344–D354.

(47) Altschul, S. F.; Gish, W.; Miller, W.; Myers, E. W.; Lipman, D. J. Basic local alignment search tool. J. Mol. Biol. 1990, 215, 403–410.

