## Supporting Information for "Screening the PRISM library against *Staphylococcus aureus* reveals a sesquiterpene lactone from *Liriodendron tulipifera* with inhibitory activity"

**CONTENTS**

**Spectroscopic data for epi-tulipinolide, tulipinolide, and honokiol**

*Epi-tulipinolide*: colorless oil; $^1$H NMR (500 MHz, $CDCl_3$) $\delta$ 6.30 (1H, d, $J$ = 3.5), 5.72 (1H, m), 5.59 (1H, d, $J$ = 3.0), 5.11 (1H, t, $J$ = 9.2), 4.87 (1H, m), 4.76 (1H, d, $J$ = 9.8), 2.90 (1H, m), 2.82 (1H, dd, $J$ = 14.6, 5.2), 2.40-2.20 (4H, ovlp), 2.09 (1H, m), 2.06 (3H, s), 1.75 (3H, s), 1.50 (3H, s). *m/z* 313 $[M+Na]^+$.

*Tulipinolide*: colorless oil; $^1$H NMR (500 MHz, $CDCl_3$) $\delta$ 6.35 (1H, m), 5.82 (1H, m), 5.03 (1H, m), 4.92 (1H, m), 4.84 (1H, m), 4.76 (1H, m), 3.03 (1H, m), 2.53 (1H, m), 2.43 (1H, m), 2.32 (1H, m), 2.21 (1H, m), 2.09 (3H, s), 1.71 (3H, s), 1.57 (ovlp). *m/z* 313 $[M+Na]^+$.

*Honokiol*: white solids; $^1$HNMR (300 MHz, $CDCl_3$) $\delta$ 7.20 (2H, s), 7.02 (2H, m), 6.90 (2H, t), 6.06 (1H, m), 5.97 (1H, m), 5.20 (2H, m), 5.05 (2H, m) 3.44 (2H, m), 3.33 (2H, m); *m/z* 267 $[M+H]^+$.

**Table & Figures**

**Table S1.** Potential sesquiterpene lactone biosynthetic enzyme amino acid sequences

**Figure S1.** Evaluation of tulip tree (*Liriodendron tulipifera*) extract against MSSA.

**Figure S2**. HPLC-DAD of the tulip tree extract. UV monitoring at 210 (black), 228 (blue), 254 (pink), and 320 (brown).

**Figure S3.** Antibacterial activity of *L. tulipifera* chromatography fractions on MSSA. Samples were prepared at 10 mg/mL and disks contained 200 μg of each fraction. Zones of inhibition were evaluated after 24 h after incubation at 37°C. Controls were (+) gentamicin sulfate (20 μg/disk) and (-) methanol (20 μL/disk).

**Figure S4.** Antibacterial activity of *L. tulipifera* HPLC fractions on MSSA.

**Figure S5.** $^1$H NMR of peak 1 (epi-tulipinolide) (500 MHz, $CDCl_3$).

**Figure S6.** $^1$H NMR of peak 2.1 (laurenobiolide) (500 MHz, DMSO-$d_6$).

**Figure S7.** HSQC of laurenobiolide (DMSO-$d_6$).

**Figure S8.** HMBC of laurenobiolide (DMSO-$d_6$).

**Figure S9.** COSY of laurenobiolide (DMSO-$d_6$).

**Figure S10.** HRESIMS of laurenobiolide (*m/z* 313.1418) recorded on SCIEX Triple TOF mass spectrometer.

**Figure S11.** $^1$H NMR of peak 2.2 (tulipinolide) (500 MHz, $CDCl_3$).

**Figure S12**. Cytotoxicity of isolated compounds from *L. tulipifera* (laurenobiolide, epi-tulipinolide, and tulipinolide) on human keratinocyte skin cells. Bars represent mean viability values compared to DMSO control (n=4) with error bars indicating standard deviation.

**Figure S13.** HPLC-DAD evaluation of epi-tulipinolide (E), laurenobiolide (L), and tulipinolide (T) in different parts of *L. tulipifera*.

**Figure S14.** HPLC-DAD evaluation of epi-tulipinolide (1), laurenobiolide (2.1), and tulipinolide (2.2) in different parts of *L. tulipifera* twigs (outer twig bark covering and inner twig material).

**Figure S15.** MSSA inhibition assay with *L. tulipifera* plant parts.

**Figure S16.** Confirmation of costunolide in *L. tulipifera*, *M. acuminata*, and *M. virginiana*.

**Figure S17.** Confirmation of dehydrocostus lactone in *M. macrophylla* and *M. virginiana*.

**Figure S18.** LC-MS/MS analysis of the *M. denudata* branch $CH_3OH$ extract.

**Figure S19.** Species identification of plant specimen (URI12) used in the current report.

**Table S1.** Potential sesquiterpene lactone biosynthetic enzyme amino acid sequences

| Annotation | Sequence |
|---|---|
| Farnesyl pyrophosphate synthase-like | MAAATSNGKSSGLRSVFLQVYARLKSELLQDPAFDWTEDSRQWIDRMLEYNVPGGKLNRGLSVIDSYKLLKDGQELSEDE<br>IFLSCSLGWCIEWLQAYFLVLDDIMDGSHTRRGQPCWFRVPKVDMIAINDGILLRNHIPRILRKNFRERPYYVDLLDLFNEV<br>EFQTASGQMLDLITTHEGEKDLSKYTMPVYCRIVQYKTAYYSFYMPVACALLMSGENLDNFTDVKNILIEMGTYFQVQDD<br>YLDCFGDPKVIGKIGTDIEDFKCSWLVVQALERADENQRKILSENYGKSDAAHVAKVKQLYKDLDLESVYLEYESKSYEKLIA<br>SIEVQPSKSVQEVLKSFLGKIYKRQK* |
| Cytochrome P450 | MLSPLSPYIAYMRYPIHLSPLALSHKISNLHLKREKIQKRGKMALFLLMIACACIYLLYLQRRKQGLPPGNLGLPFIGETLQLVS<br>AYKTDNPEPFIDARVRRYGSLFTTHVFGEPTVFSTDPEANRFVLQNEGKLFESSYPSSISNLLGRHSLLLMKGNLHKRMHSLT<br>MSFANSSIIRDHLLVDIDRLVRFNLQRWDGLILLQDETKKITFELTVKQLMSFDPGEWTESLRKEYLLLIEGFFSVPIPFFFTTY<br>GRALQARTKVAAALRERVRERKERRRNRKGEEQKDMLGALLDEGEGGFSEEEAVDFLLALLVAGYETTSTIMTLAVKFLTE<br>TPSALALLKEEHEGIRAKKKESEALDWSDYKSMPFTQCVSLLFPLIN* |
| Cytochrome P450, E-class, group IV | MLSPLSPYIAYMRYPIHLSPLALSHKISNLHLKREKIQKRGKMALFLLMIACACIYLLYLQRRKQGLPPGNLGLPFIGETLQLVS<br>AYKTDNPEPFIDARVRRYGSLFTTHVFGEPTVFSTDPEANRFVLQNEGKLFESSYPSSISNLLGRHSLLLMKGNLHKRMHSLT<br>MSFANSSIIRDHLLVDIDRLVRFNLQRWDGLILLQDETKKITFELTVKQLMSFDPGEWTESLRKEYLLLIEGFFSVPIPFFFTTY<br>GRALQARTKVAAALRERVRERKERRRNRKGEEQKDMLGALLDEGEGGFSEEEAVDFLLALLVAGYETTSTIMTLAVKFLTE<br>TPSALALLKEEHEGIRAKKKESEALDWSDYKSMPFTQCVINETLRVANIISGVFRRAVSDVNIKGYTIPKGWKVFASFRAVHL<br>DQDLYKDARTFNPWRWQVLCLLFPFTEGTGIVPDMLCFNPLAHAL* |
| acetyl-CoA acetyltransferase, cytosolic | MAPAAASDSIKPRDVCVVGIARTPMGGFLGTLSSLSATKLGSIAIECALKRADIDPKLVQEVYFGNVLSANLGQAPARQAAL<br>GAGIPNTVICTTINKVCASGMKATMLAAQSIQLGINDVVVAGGMESMSNAPKYLSEARKGSRLGHDTIVDGMLKDGLW<br>DVYNDYGMGMCAELCADQHSITREEQDSYAIQSFELGIAARNGGAFAWEIVPVEVSGGRGKPSVLVDKDEGLEKFDPVK<br>LRKLRPNFKENGGSVTAGNASSISDGAAALVLVSGEKALELGLQVIAKISGYADAAQAPELFTTAPALAIPKAISNAGLEASQI<br>DYYEINEAFAVVSVANQKLLGIHPDKLNVHGGAVSLGHPLGCSGARILVTLLGVLRQRNGKYGVAGICNGGGGASALVLEL<br>MPVIRAERSSL* |
| terpene cyclase | DLSFQLHLSVEMAHQGPPSLFSNLQATEIPKPGVIRPTAGFHPTAWGDHFLNYSGENKNVDAWTKKVEMLKEEVRRMLV<br>NNKVPVQEMNLIDDIQRLGVAYHFEKEIDKALQHIYDEYQNVHYDDLYVVALRFRLLRQGGYNVSSDVFSKFKGEDGNFK<br>ATLSRDVKGMLSLYEAAYFSIQGEDILDEAIVFTSGHLTTIMAHLHRPLAENARRALELPLHKRIPRLDARYYISLYEEDKSHN<br>DVLLELARLDFNLLQLLHQRELRDLSRWWKDLGLATKYPFARDRLVETYFWILGVYFEPQYTRARAIITKIMKIASIIDDIYDA<br>YATFDELEIFTNAVLRWELEAAEELPEYMKACYLALLNTIDEIESQMMPDEKFYRTNFIKREMKVLVHSYLDEAKWVKRRYV<br>PTFGEHLDVSLLSCGFPLLTGVAYVGMGDLASKEVFDWLNTHPKFVMDLSIVCRLVDDIAGHHFEQEREHVASTVECYMK<br>EHGVSEQEACTKLREMIATAWKDVNKACLRPTVIPLPLLVRGANSTRVIEDLYIRGDGYTHSKYETKERVTVVLVDPVPIPL* |
| terpene cyclase | MAHQGPPSPYSTLQATEIKKPEVVRPTAGFHPSVWGDRFLDYSEEQKVVDEWTGKVEVLQEEVRWMLINNKGSVQEM<br>NLIDDIQRLGVAYHFEKEIDEALHRIYDAYTNVHYDDLYAGALRFRLLRQSGYNVSSDVFSKFKDEDGNFKETLSSDVRGML<br>SLYEAAYLGIQGEDILDEAIVFTSGHLKTIVAQLHHPLVEKVRHALVLPLHKRVPRLEARYYISLYQEEESHNDVLLELARLDFN<br>LLQSLHQRELRDLSRWWKDLGLVTKYPYARDRLVEAYYWILGVYFEPQYSRARVILTKIFKLTSIIDDTYDAYATLDEVQIFTN<br>AIHRWELEAAEGLPDYMRACYLALLRTVDEIEDQMMADEKFYRTNWLKREMKVLVQAYFDEAKWMNSGHVPTLKEHL<br>DVSLISAGYIFVYGVAFVGMGDEASKEIFDWMTAYPKFIMDLSIIARVGDDIGGHKFEQEREHVASTVECYMKEHGVSDKE<br>ACIKLQEMITTAWKDLNKACLRPTVIPLPLLLRGLNLARVMEELYKQGDGYTHSNNETKEKIMAVLVDPILIRG* |
| terpene cyclase | CLEKTHSACLLLLSSFHSIMALILGGGHSDGPTNQGKNGKKEIGRASANYHPSVWEDRFIAASPDDKELDPYTKQRADMLK<br>EEVKKMLCKVNNSVQKLNSIDAIQRLGVAYHFETDIEKELHRMYDGYNDDGDNLHVVALRFRLLRQHGYNVSSDVFRKFK<br>DNKGKFKATMSSDIRGLLSLYEAAYLSIHGDDILDEAITFTTMHLKSAMLHLTSSLAKLVELALEVPLRKCVERLQSRYYISIYE<br>EEKERSDILLEFAKLDFNLLQSLHRSELRDISRWWKENDFAVKLPFIRDRVVECYFWILGVYFEPHYSRARHMMTTIISLTSI<br>MDDIYDVHGTLEELELYTAALESWDRGAVDRLPEYMKVHFVALLDAVDGFEDELSQEGKSYRISYLKEVYKVLARAYLQEA<br>RWASSEYVPTYNEYMEVAQISSAYPLLTVISLVGMGDIVTKEAFEWAVNSPKVVTACSVICRLKDDITSSELEQQRSHVASA<br>MQCYMREHSASYTDTCEKFEEMVAIAWKEVNKECLKPTHVPMSVIMRTVNLARVIEFLYGHQDGYTNSTRETKERIQMV<br>MVDPVPV* |
| terpene cyclase | MALILGNGHSDIPTKNQVETKGRKEIGRGCANYHPSVWGDQFVTLSPDEMKIDVQTKQRAEILKEELKRMLLNVSDSLQE<br>LTLINEIQRLGVAYHFEKEIKDALYRMYDAHSNGGNDVSDDLHAVALWFRLLRQQGYNVSSNVFRRFKDENGEFKATLKD |

| | |
|---|---|
| | DIRGLLSLYEAAYLGTREDNILDEAINFTTEQLKSAMSHLSSPLSTLVQLALAVPLHRRVERLQSRYYISIYQQEKERNDVLLEF<br>AKLDFNMLQSLHKKELSDISRWWKENDFSRKLPFIRDRIVELYFWVLEVYFEPQYARARRMMTTIISLTSILDDIYDVYGTLE<br>ELELYTVAIESWDWAAMDQLPDYIKPHYTALLNAVEKFEDELSQEGKSYRIPYLKKALTVLAKGYLEEARWTSAEHTPTLEE<br>YMKIALITNGYPMLTIASMVGMGDIVTKEAFEWAINVPKVVEASAAICRLRDDITSNEFEQERTHVASGIQVYMKEYNTTY<br>EEACNIFLQKTANAWKDANMECMEPTPVPREVIKRPINLGRVIELLYQHKDSYTNSAFETKEHITMVLVDPIPL* |
| terpene cyclase | RRKLRSSREAVPKDHTRHAYTLSSLPPFISSFHSIMALIFGSGHSYSPTTQGKNGKKEIGRSCANYHPSVWGDSFIATSPHDK<br>ELDPSTMRRVEKLKEEIKKMLCDVDNLVEKLNLIDAIQRLGIAYHFETDIEKELHKVYDGYNDNGDNLHVIALQFRLVRQQG<br>YNVSSDVFRKFKDNEGKFKAKLSSDIRGLLSLYEAAYLSTHGDDILDEAIIFTSEHLKSALPHLTSPLTKLVQLALEVPLWRRVE<br>RLQSRYYISIYEEEEERSDVLLEFAKLDFNLLQSLHRRELRDISKWWKKNDFAAKLPFIRDRVVECYFWILGVYFEPHYSRARR<br>MMTTIISLTSIMDDIYDVYGTLEELELYTSVIESWDRGAVDKLPEYMKGHFVALLDAVDGFEDELSREGKSYRISYLKEAYNG<br>VARAYLQEARWASSEYVPTYEEYMEVAQISSAYPMLIVISQVGMGDIVTKEALEWAINIPKVVTACSVICRLKDDITSSKLE<br>QARGHVASAMQCYMREHGNSYTDTCEKFQEMVAMAWKEVNKECLKPTHVPMPVIMRAVNLARVIELLYVHQDGYTN<br>STCETKERIAMVMVDPLVV* |
| terpene cyclase | SFPTRRSSDLFQLHPSIEMAHQGPPSLCSTLQANKIRKSEVVRQTAGFHPTVWGDHFLNYSVEDKNVDAWTRKVEVLKEE<br>VRKMLVNAKGSVQEMILINDIQRLGVAYQFEKEIDEALSSIYDAYTNVHYDDLLAVALRFQLLREAGFNVSSDVFRKFKDDD<br>GNFKATLSSDVRGMLCLYEAAYFGIQGEEILDEAIVFTSGHLNSIMPHLHHPLVAKVQRALELPMRKRILRLEARYYISLYQD<br>EESHNDVLLELARLDFNILQSLHQTELKDLCRWWKDLGLATKYPFARDRLVEGYFWVLGVYFEPQYTCARAILTKIFKILSIM<br>DDIYDTYATLDELETFTNAIHRWELEAAEGLPDYMKACYLALLNTLNEIEGQMMPDEKVYRTNYIKREMKAMVQGYLDEA<br>KWANKRHVPTLGEHLDVSLVTAGNRLLVGVTYAGMGDLASKEVFDWLNTHPKFIMDLNIIGRLVDDIVGHQFEQERMH<br>AASTVECYMKDHGVSEQEACAKLQEMVATAWKDLNKACLRPTVIPLPLLLPAIGLVRVVEDLYIHGDGYTDSRNETKEKVI<br>MVLVDPIPIPMRK* |
| terpene cyclase | MTTCFIVSPEAMKRCNFRFQTDAIFDLILFLYMMLMQRLNSHKRRGEELKEVVRNMLCTIDDPVLKMNLIDAIQRLGVAY<br>HFEMDIDKALRRMYDDNINGNDDGFDLQALALQFRLLRQQGYNVSSSVFTKFKDDEGNFNAILSSDTRSLLSLYEAAFLGI<br>HGDDILDEAITFTTAHLKSTLSHLTPPLKKLVELALEIPLQRCFERLQTRYYISIYEEDNERNDVLLEFAKLEFHIFQSLHQRELR<br>DMSLWWKEMNLIAKLPFARDRVVEGYFWTVGVYFEPHYSLARMIMAKMIALTTVMDDIYDIYGTLEELELLTATIQRWD<br>RGDMDQLPDSMKVFFIALLDTVDAFEDELTREEKSYRMYYLKEAIKDQAKVYLLEARWASSGYVPTSEEYMKVAVISAAYP<br>MLFVAFLIGMGEVVTKEVLEWAKHVPMMMRTCSTMVRLMDDIQSSKLERERQHVSSAVECYMKEHGSSYQETIQKLRE<br>MVASGWKDINKECLKPTPAPTAVINVILNFTRVLELIYRYRDGYTDSTVETKEQIALVLVDPVPL* |

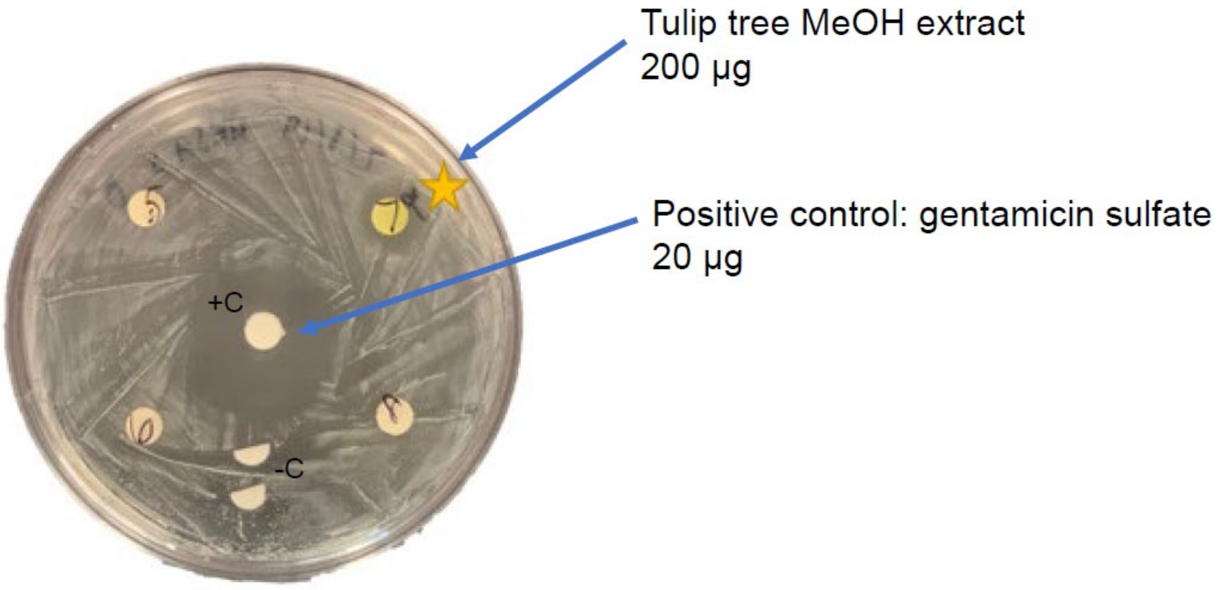


**Figure S1.** Evaluation of tulip tree (*Liriodendron tulipifera*) extract against MSSA. A zone of inhibition was observed around the disk containing the tulip tree extract.

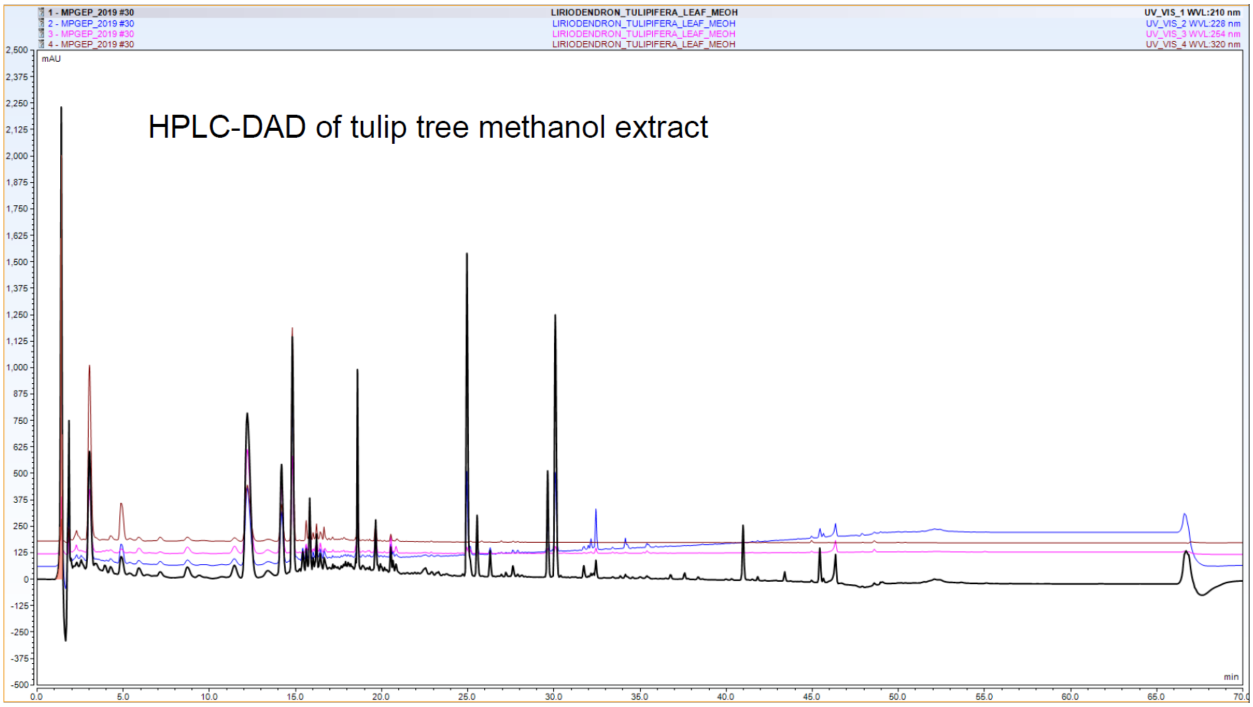


**Figure 2**. HPLC-DAD of the tulip tree extract. UV monitoring at 210 (black), 228 (blue), 254 (pink), and 320 (brown).

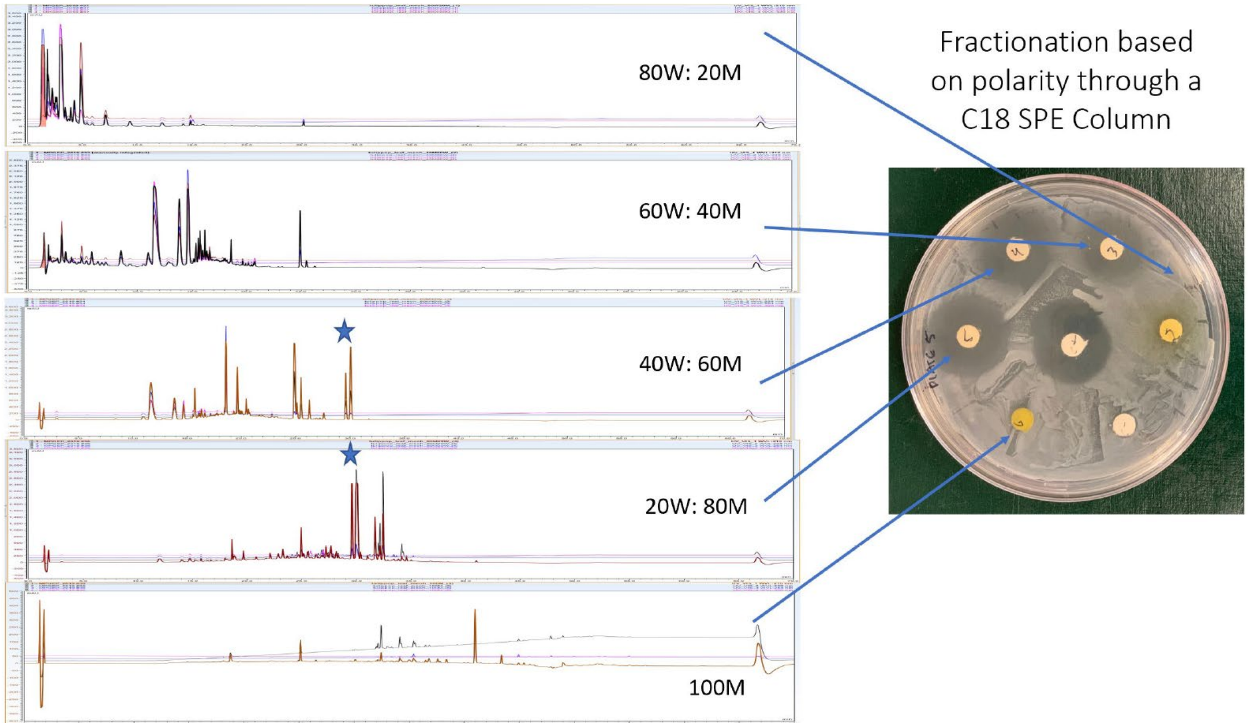


**Figure S3.** Antibacterial activity of *L. tulipifera* chromatography fractions on methicillin-susceptible *S. aureus.* Samples were prepared at 10 mg/mL and disks contained 200 µg of each fraction. Zones of inhibition were evaluated after 24 h after incubation at 37°C. Controls were (+) gentamicin sulfate (center) and (-) blank. Stars indicate common peaks of interest from the most active fractions.

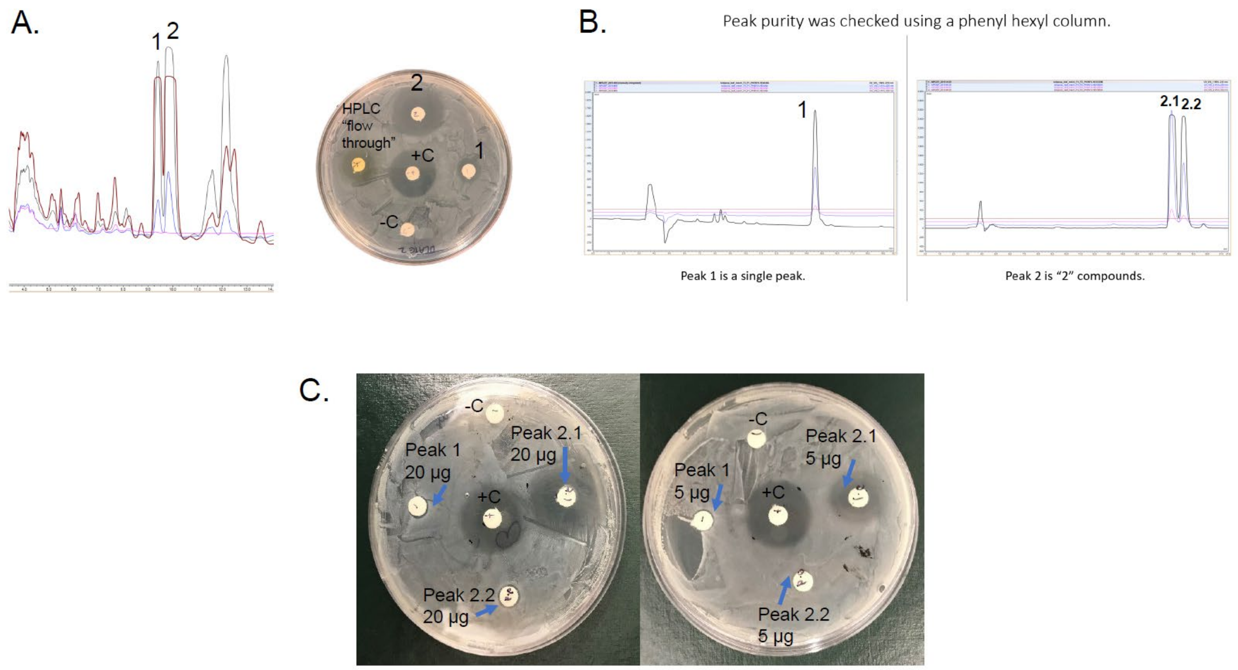


**Figure S4.** Antibacterial activity of *L. tulipifera* HPLC fractions on methicillin-susceptible *S. aureus.* A) Initial peaks collected and tested (1 and 2) with corresponding MSSA test results. B) Peak 2 was composed of two compounds following HPLC analysis using phenyl hexyl column (250 x 10 mm, 5 µm). C) Peak 2.1 (laurenobiolide), 2.2 (tulipinolide), and 1 (epi-tulipinolide) tested at 20 and 5 µg, respectively. Only 2.1 showed zones of inhibition. Controls were (+) gentamicin sulfate (20 µg) and (-) blank.

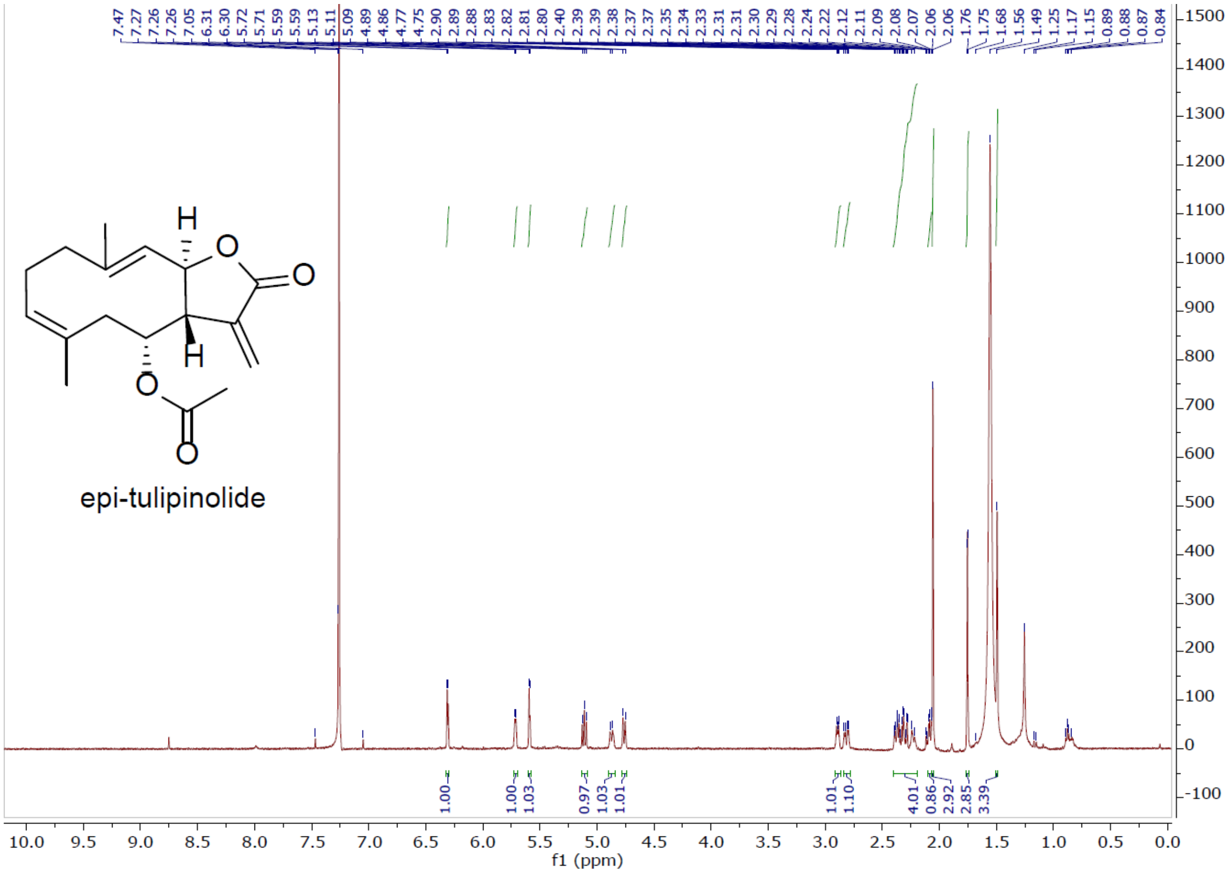


**Figure S5.** $^{1}$H NMR of peak 1 (epi-tulipinolide) (500 MHz, $CDCl_3$).

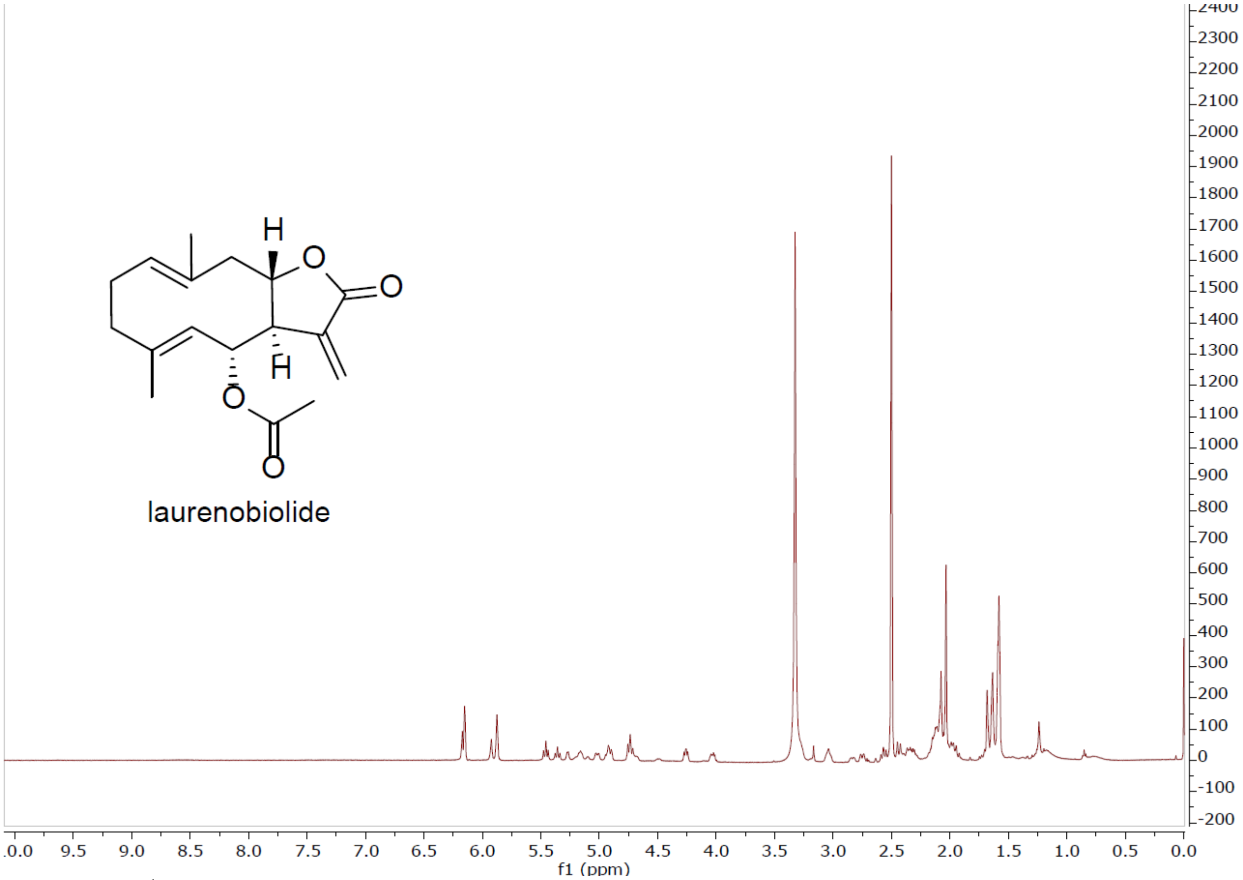


**Figure S6.** $^{1}H$ NMR of peak 2.1 (laurenobiolide) (500 MHz, DMSO-$d_6$).

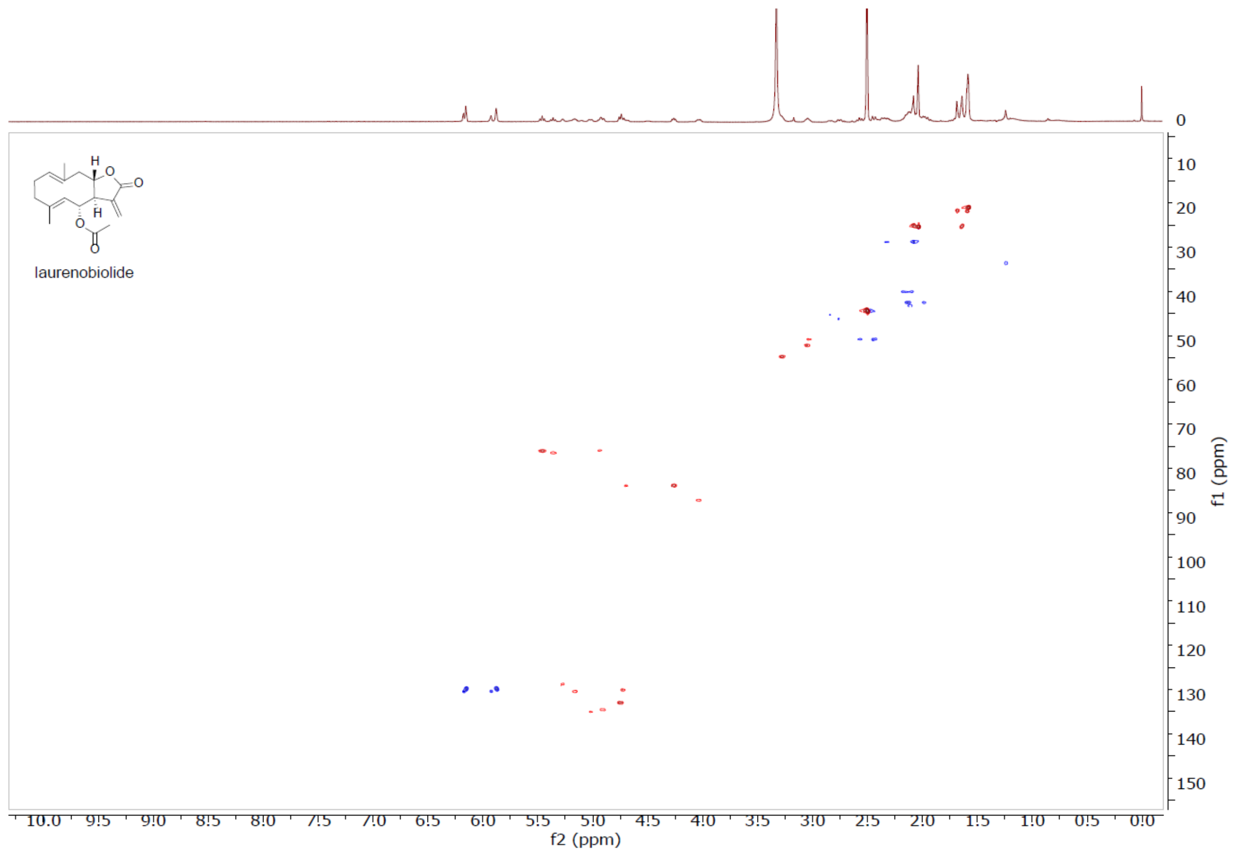


**Figure S7.** HSQC of laurenobiolide (DMSO-$d_6$).

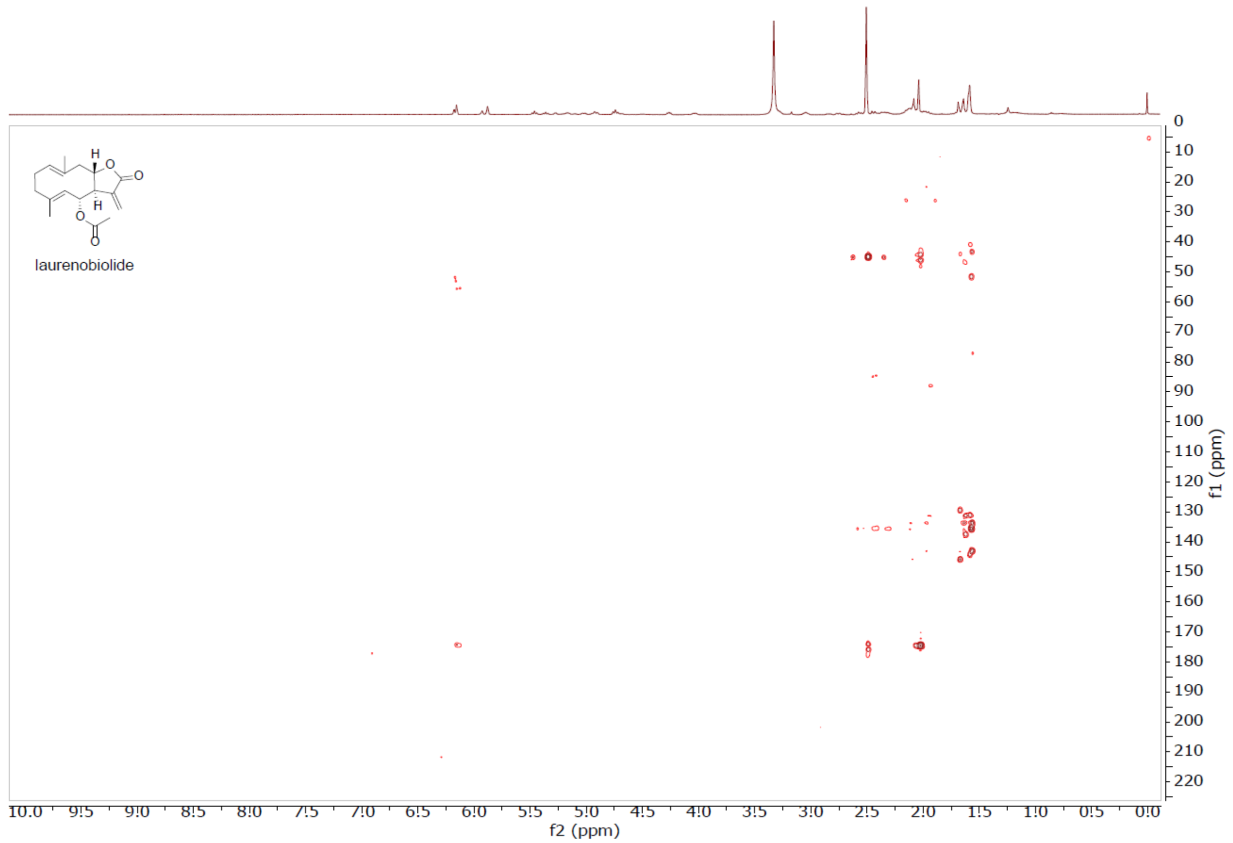


**Figure S8.** HMBC of laurenobiolide (DMSO-$d_6$).

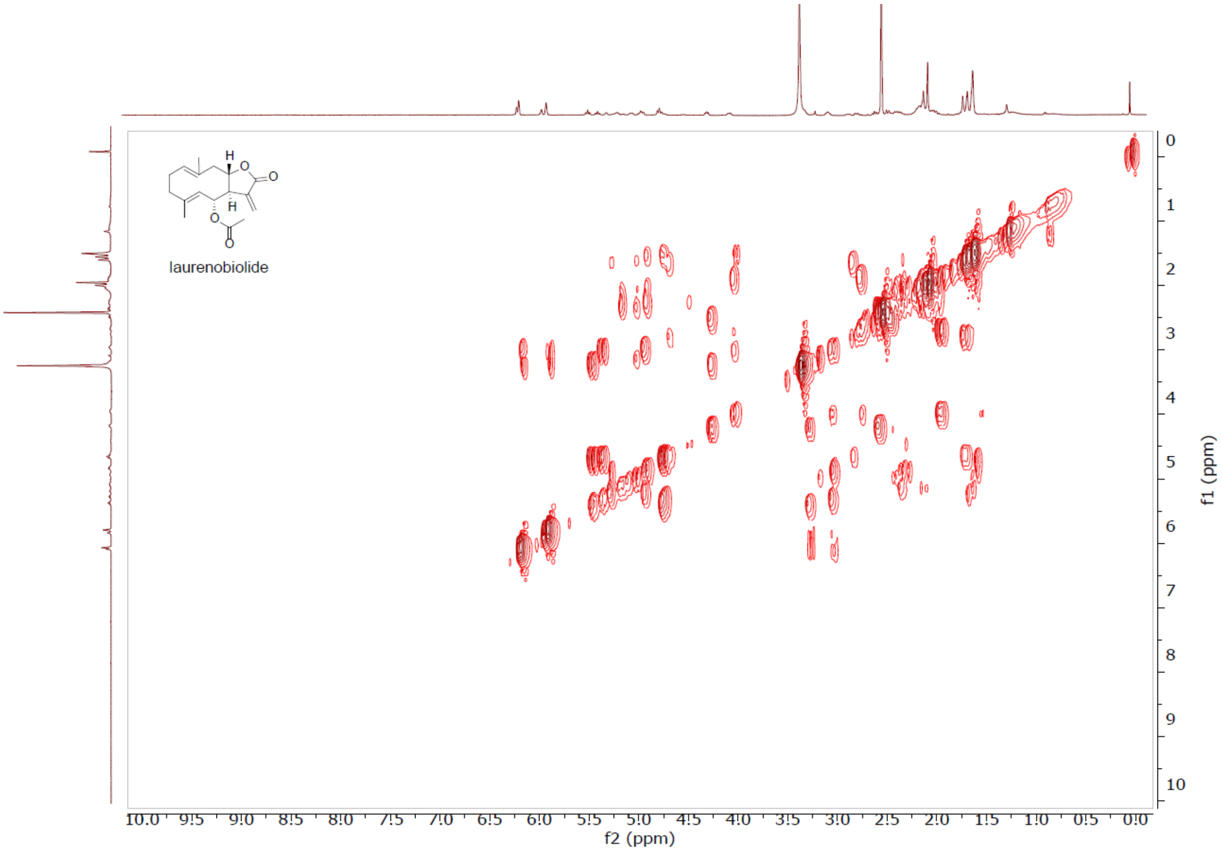


**Figure S9.** COSY of laurenobiolide (DMSO-$d_6$).

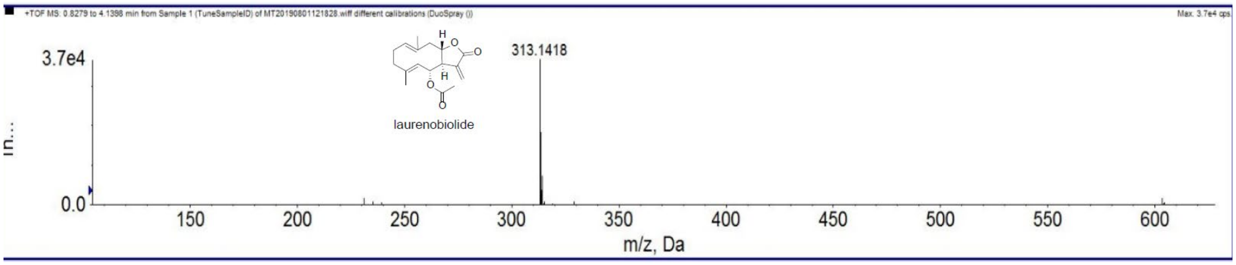


**Figure S10.** HRESIMS of laurenobiolide (*m/z* 313.1418) recorded on SCIEX Triple TOF mass spectrometer.

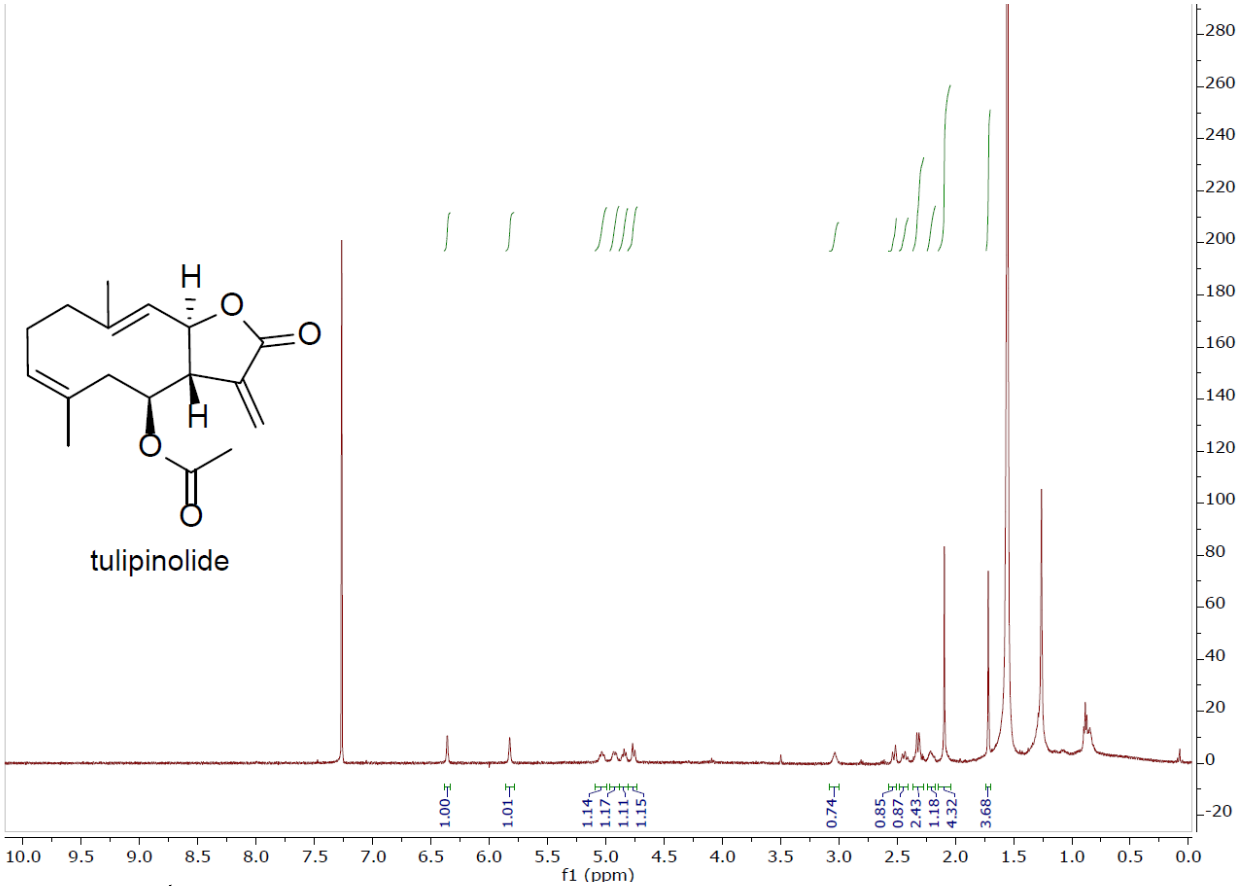


**Figure S11.** $^{1}$H NMR of peak 2.2 (tulipinolide) (500 MHz, $CDCl_3$).

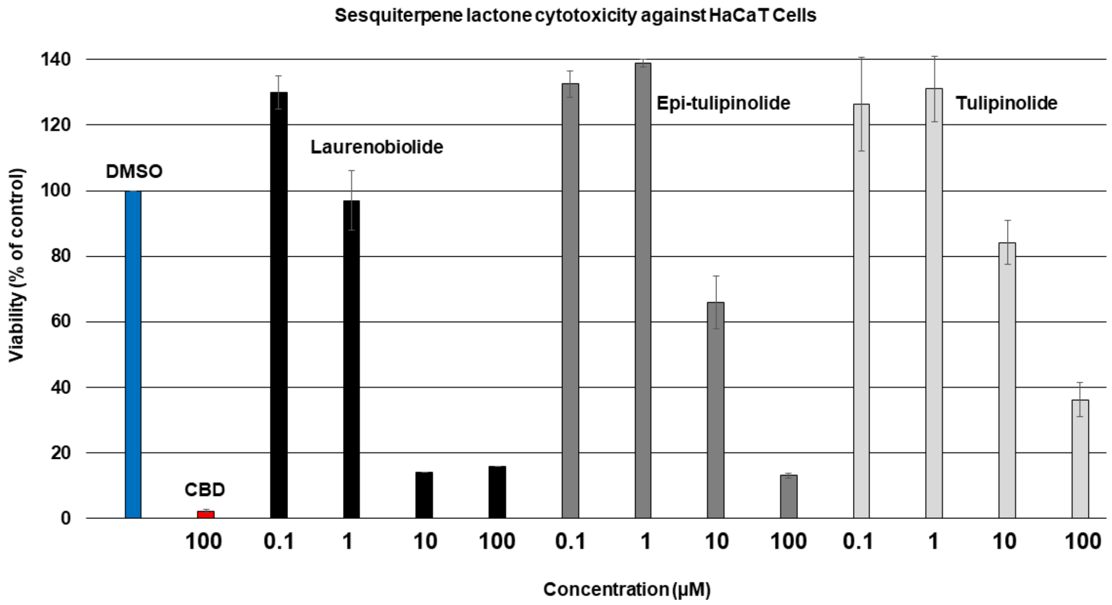


**Figure S12**. Cytotoxicity of isolated compounds from *L. tulipifera* (laurenobiolide – black bars, epi-tulipinolide – dark gray bars, and tulipinolide – light gray bars) on human keratinocyte skin cells. Bars represent mean viability values compared to DMSO control (blue bar) (n=4) with error bars indicating standard deviation.

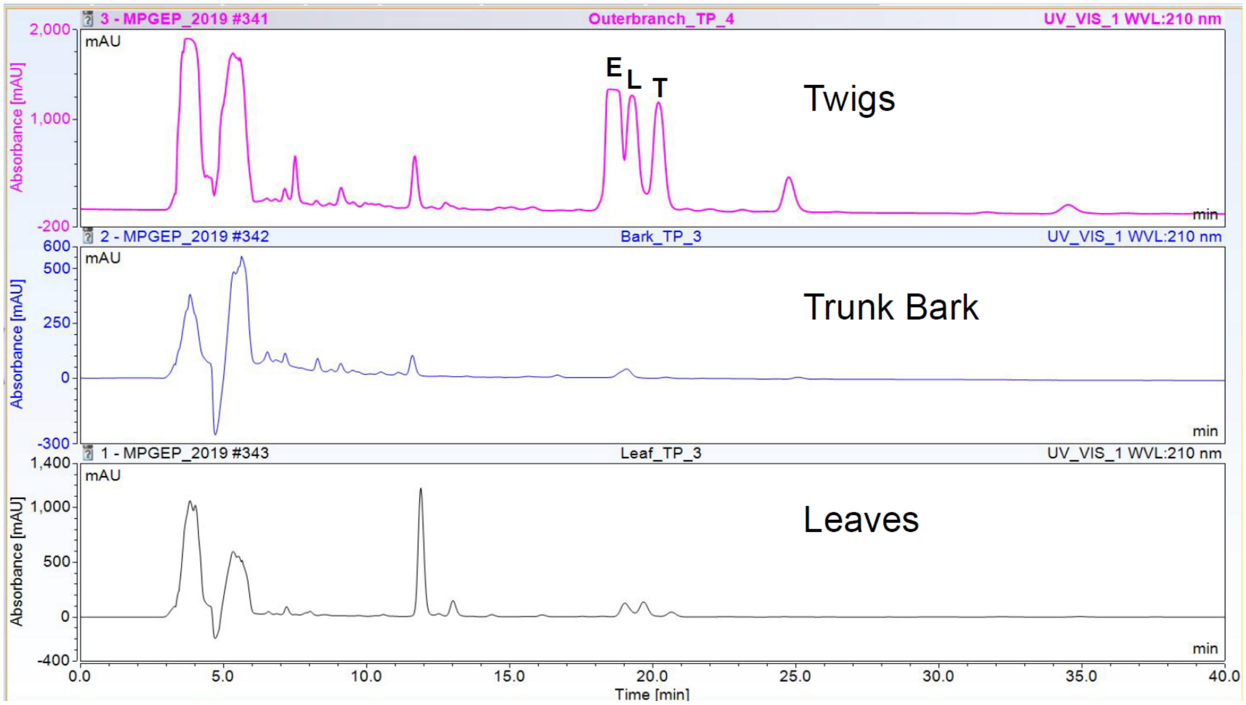


**Figure S13.** HPLC-DAD evaluation of epi-tulipinolide (1), laurenobiolide (2.1), and tulipinolide (2.2) in different parts of *L. tulipifera*.

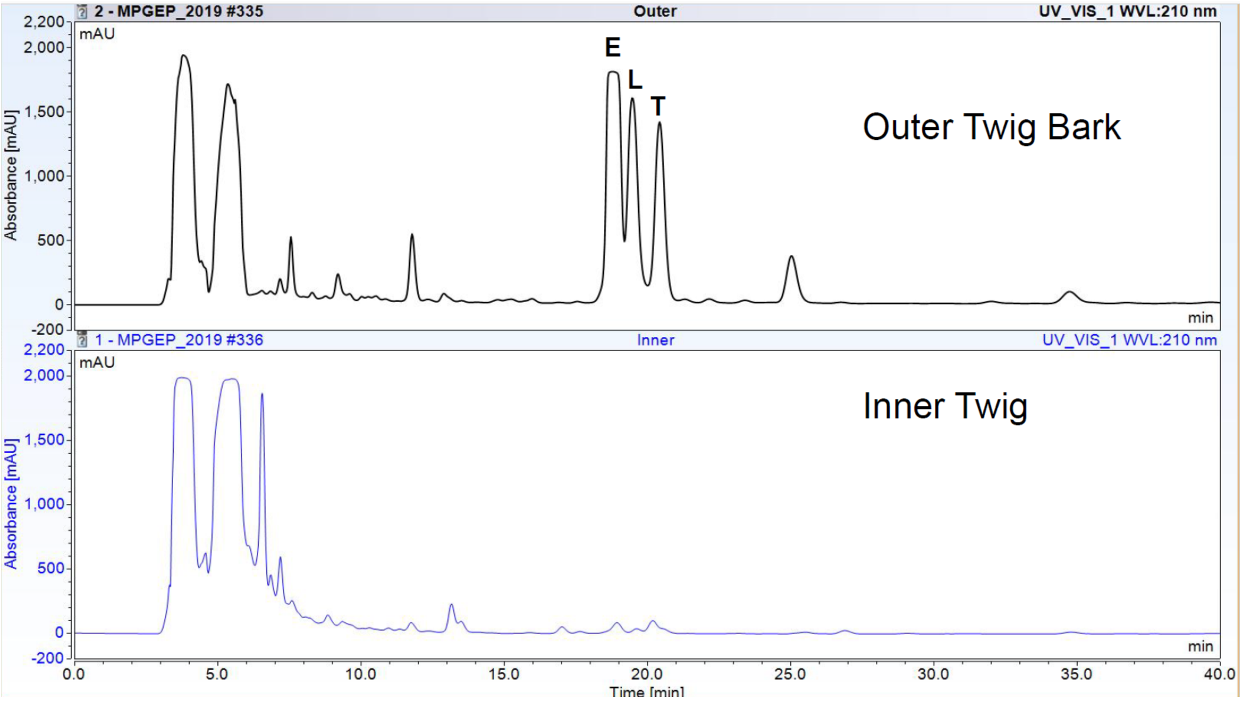


**Figure S14.** HPLC-DAD evaluation of epi-tulipinolide (1), laurenobiolide (2.1), and tulipinolide (2.2) in different parts of *L. tulipifera* twigs (outer twig bark covering and inner twig material).

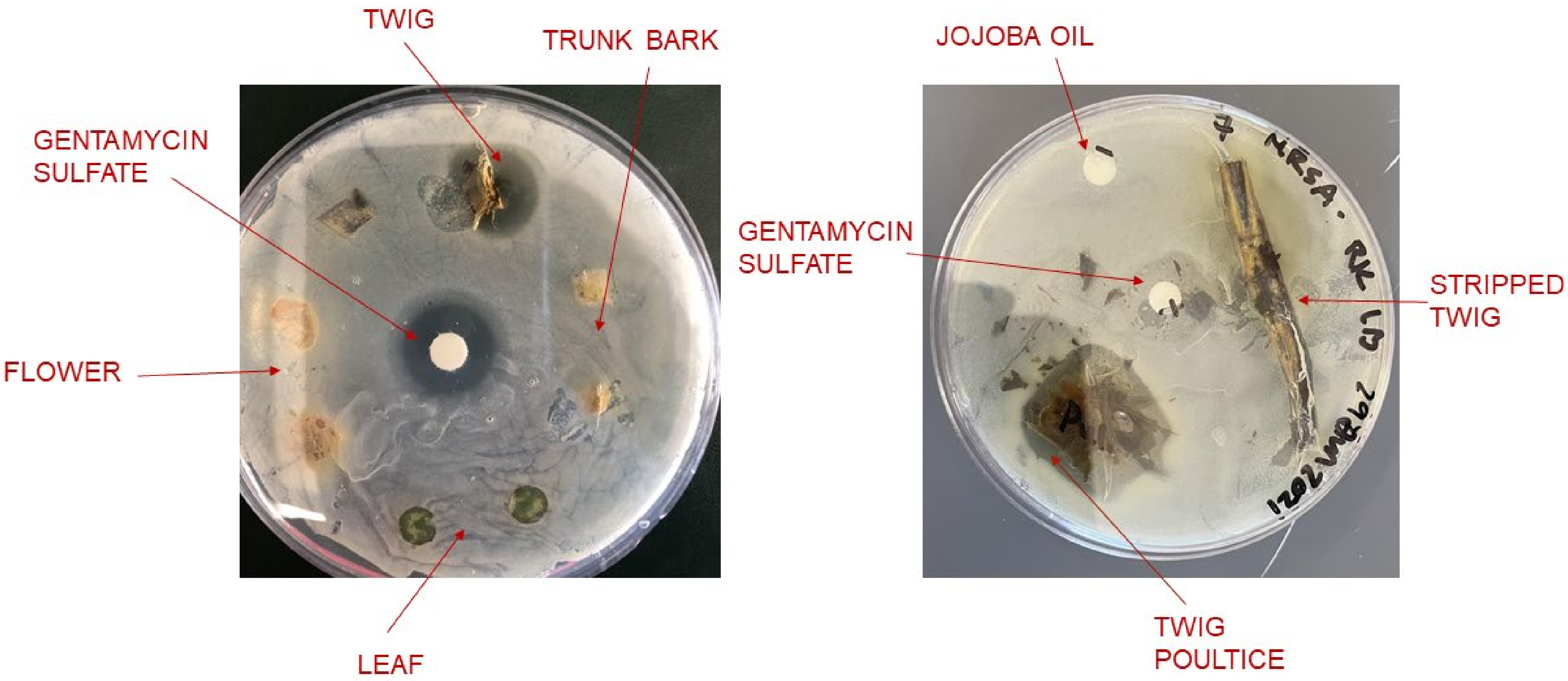


**Figure S15.** MSSA inhibition assay with *L. tulipifera* plant parts. Left panel shows twigs, trunk bark, flower, and leaf directly placed onto MSSA plates. Plant parts were surface sterilized with methanol and dried prior to evaluation. The inhibition can be seen for the twig specimen. Right panel shows the inhibitory effect of a twig poultice using jojoba oil as the carrier while a stripped twig without outer bark does not show any inhibitory effects. Controls were (+) gentamicin sulfate and (-) jojoba oil.

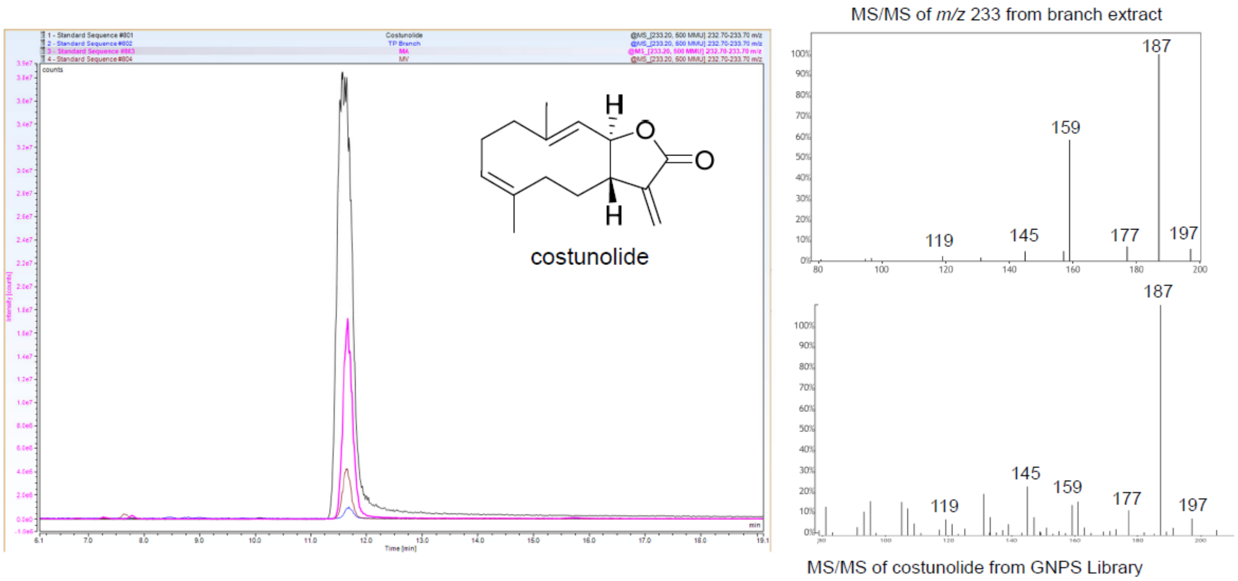


**Figure S16.** Confirmation of costunolide in *L. tulipifera*, *M. acuminata*, and *M. virginiana*. Constunolide standard is in black, while *L. tulipifera*, *M. acuminata*, and *M. virginiana* are in blue, pink, and brown, respectively. MS/MS fragmentation comparison of *m/z* 233 from the *L. tulipifera* extract with the MS/MS spectrum of constunolide available in the GNPS library is shown at right.

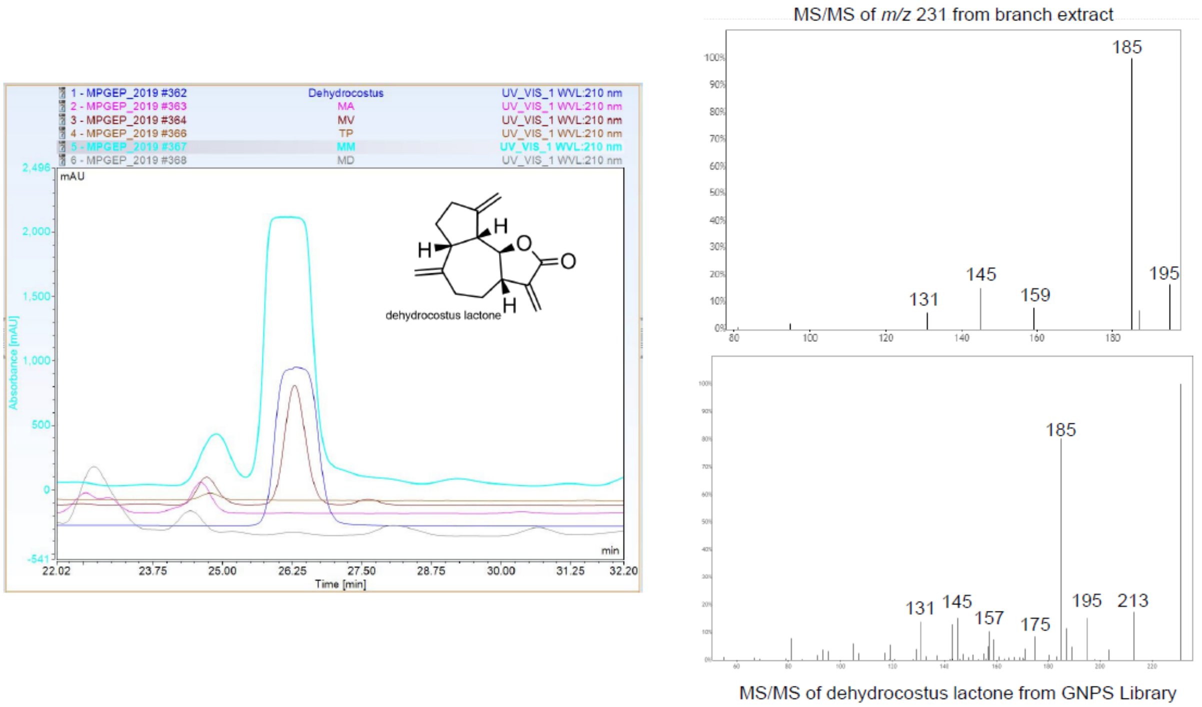


**Figure S17.** Confirmation of dehydrocostus lactone (blue) in *M. virginiana* (brown) and *M. macrophylla* (cyan).

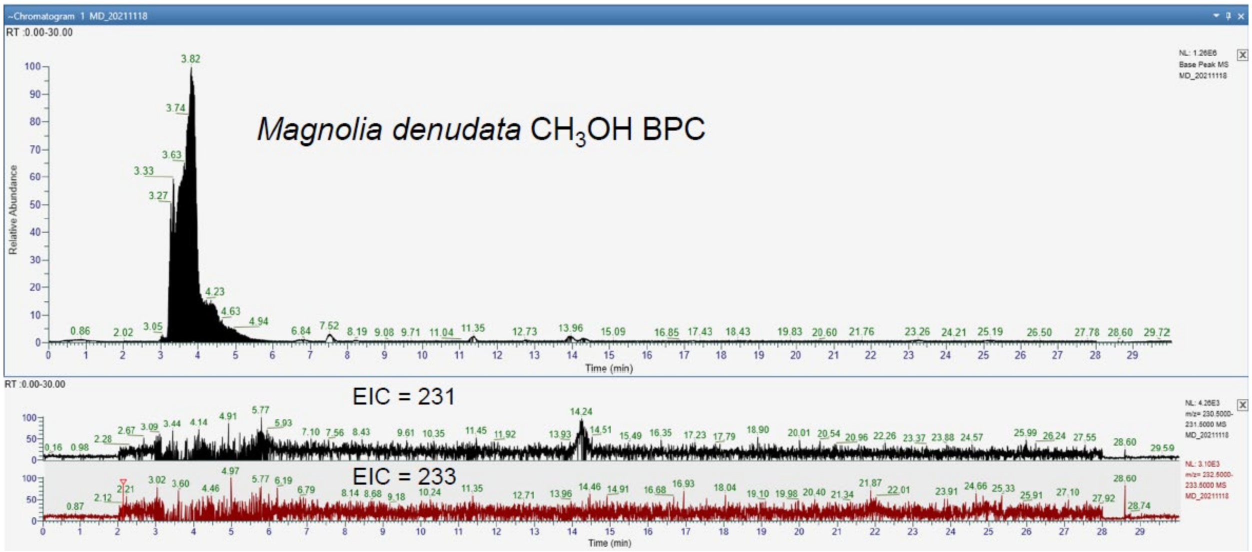


**Figure S18.** LC-MS/MS analysis of the *M. denudata* branch $CH_3OH$ extract. The top panel is the base peak chromatogram. The middle panel shows an EIC for *m/z* 231 and the bottom panel shows an EIC for *m/z* 233.

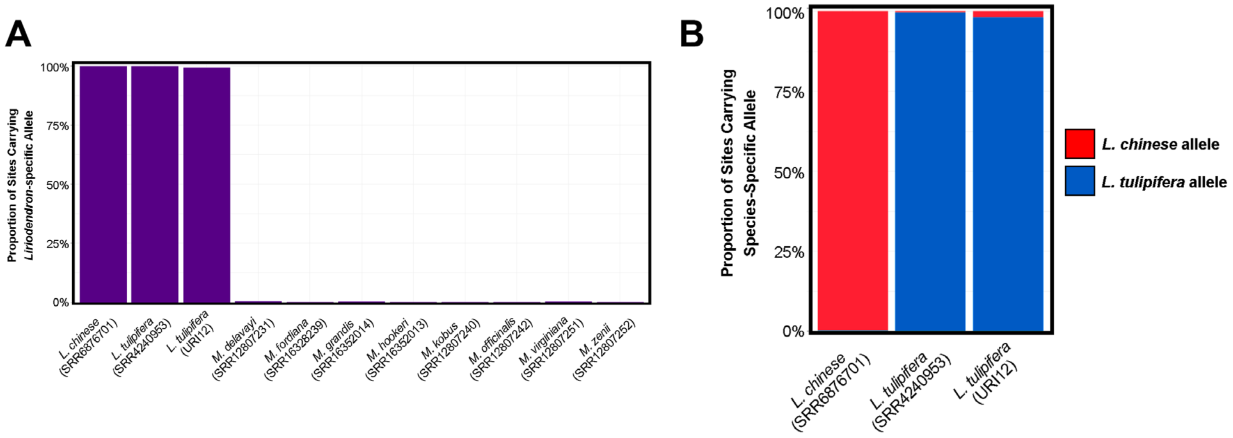


**Figure S19.** Species identification of plant specimen (UIR12) used in the current report. Proportion of sites carrying alleles matching either (A) the *Liriodendron* genus-specific alleles or (B) the *L. tulipifera* or *L. chinese* specific alleles was tabulated. This SNP analysis clearly positioned URI12 as *Liriodendron tulipifera*.
